# Activating a collaborative innate-adaptive immune response to control breast and ovarian cancer metastasis

**DOI:** 10.1101/2020.07.13.200477

**Authors:** Lijuan Sun, Tim Kees, Ana Santos Almeida, Bodu Liu, Xue-Yan He, David Ng, David Spector, Iain A. McNeish, Phyllis Gimotty, Sylvia Adams, Mikala Egeblad

**Affiliations:** Cold Spring Harbor Laboratory, Cold Spring Harbor, NY 11724, USA; Department of Surgery and Cancer, Imperial College London, London W12 0NN, UK; Department of Biostatistics, Epidemiology & Informatics, University of Pennsylvania, Philadelphia, PA 19104-6021, USA; Perlmutter Cancer Center, New York University, New York, NY 10016, USA

## Abstract

Many cancers recruit monocytes/macrophages and polarize them into tumor-associated macrophages (TAMs). TAMs promote tumor growth and metastasis and inhibit cytotoxic T cells. Yet, macrophages can also kill cancer cells after polarization by *e.g.*, lipopolysaccharide (LPS, a bacteria-derived toll-like receptor 4 [TLR4] agonist) and interferon gamma (IFNγ). They do so via nitric oxide (NO), generated by inducible NO synthase (iNOS). Altering the polarization of macrophages could therefore be a strategy for controlling cancer. Here, we show that monophosphoryl lipid A (MPLA, a derivative of LPS) with IFNγ activated macrophages isolated from metastatic pleural effusions of breast cancer patients to kill the corresponding patients’ cancer cells *in vitro*. Importantly, intratumoral injection of MPLA with IFNγ not only controlled local tumor growth but also reduced metastasis in mouse models of luminal and triple negative breast cancers. Furthermore, intraperitoneal administration of MPLA with IFNγ reprogrammed peritoneal macrophages, suppressed metastasis, and enhanced the response to chemotherapy in the ID8-p53^−/−^ ovarian carcinoma mouse model. The combined MPLA+IFNγ treatment reprogrammed the immunosuppressive microenvironment to be immunostimulatory by recruiting leukocytes, stimulating type I interferon signaling, decreasing tumor-associated (CD206^+^) macrophages, increasing tumoricidal (iNOS^+^) macrophages, and activating cytotoxic T cells through macrophage-secreted interleukin 12 (IL-12) and tumor necrosis factor α (TNFα). Both macrophages and T cells were critical for the anti-metastatic effects of MPLA+IFNγ. MPLA and IFNγ are already used individually in clinical practice, so our strategy to engage the anti-tumor immune response, which requires no knowledge of unique tumor antigens, may be ready for near-future clinical testing.

## Introduction

Cancer elicits an immune response, but solid tumors evade this response, in part by establishing an immunosuppressive tumor microenvironment. The tumor microenvironment is composed of tissue-resident cells (*e.g.*, fibroblasts and endothelial cells), innate immune cells (*e.g.*, macrophages), and adaptive immune cells (*e.g.*, T cells) (*1–3*). It was discovered over a century ago that intratumoral injection of dead bacteria, named “Coley’s toxin” after the physician who devised the treatment, led to durable anti-tumor responses in some patients (*4*). It is likely that the tumor responses were caused by the activation of immune cells, particular macrophages, by bacterial components. However, tumor-associated macrophages (TAMs) are usually associated with pro-tumor features: TAMs promote tumor growth and metastasis directly (*5–7*). They also contribute to the tumor immunosuppressive microenvironment, allowing tumors to evade the intrinsic anti-tumor immune response, as well as T cell-based therapies, including checkpoint blockade (*8–10*).

Macrophages can take on a continuum of phenotypes between two major subtypes: classically activated and alternatively activated macrophages (*11*). Classically activated macrophages are stimulated during acute inflammation by interferon γ (IFNγ) and toll-like receptor (TLR) ligands, including bacterially derived lipopolysaccharide (LPS). These macrophages secrete pro-inflammatory cytokines (tumor necrosis factor α [TNFα] and interleukins IL-1β, IL-2, IL-6, IL-12, and IL-23) and metabolize arginine to the “killer” molecule nitric oxide (NO) through upregulation of inducible nitric oxide synthase (iNOS). Stimulating macrophages with IL-4/IL-13 leads to the alternatively activated macrophage phenotype and the subsequent production of anti-inflammatory cytokines (IL-10, TGFβ). TAMs usually exhibit traits of the alternatively activated macrophage subtype and have been clinically targeted by ablating strategies, such as anti-CSF1R therapeutics, which unfortunately have long-term toxicity and high tumor recurrence limiting their use (*12–14*). However, depending on context, macrophages in tumors can also eliminate tumor cells with NO, reactive oxygen species, or by phagocytosis, and they can stimulate cytotoxic T lymphocytes to kill tumors by presenting antigens and producing pro-inflammatory cytokines (*11, 15*). Since macrophages exhibit a high degree of plasticity, a better strategy than macrophage ablation may be to “reprogram” the TAMs to kill cancer cells and support the elimination of tumors by cytotoxic T lymphocytes (*16, 17*). Class IIa histone deacetylase (HDAC) inhibitors can reprogram TAMs to anti-tumor macrophages and increase the efficacy of chemotherapy and immunotherapy in preclinical models of breast cancer (*18*). Similarly, selective inactivation of macrophage PI3Kγ promotes an immunostimulatory transcriptional program in turn restoring CD8^+^ T cell activation and cytotoxicity and increasing survival in mouse models of head and neck squamous cell carcinoma (*19*).

Reprogramming of macrophages and/or direct stimulation of adaptive immunity have been achieved using *e.g.*, TLR agonists. TLRs are innate immunity pattern recognition receptors that play fundamental roles in activating the innate immune response (*20*). Nanoparticles loaded with R848 (a TLR7/8 agonist) effectively accumulate in TAMs, repolarize them towards an inflammatory phenotype, and show profound tumor-suppressing effects in a colon cancer mouse model (*21*). Another delivery approach besides nanoparticles is direct intratumoral injection: injection with a CpG oligonucleotide (a TLR9 agonist) reverts resistance to PD-1 blockade by expanding multifunctional CD8^+^ T cells in breast and colon cancer mouse models (*22*). In a phase I clinical study, the combination of motolimod (a TLR8 agonist, subcutaneously administered) with the EGFR inhibitor cetuximab (intravenously injected) increased secretion of IL-6, CCL2, and CCL4 and increased natural killer (NK) cell circulation and activation (*23*). Similarly, the TLR7 ligand imiquimod applied as a cream to breast cancer skin metastases showed anti-tumor activity and triggered a strong T-helper-1 (Th-1)/cytotoxic immune response in responding patients (*24, 25*). However, the therapeutic efficacy of motolimod and imiquimod is low: only two out of eleven squamous cell carcinoma patients and two out of ten breast cancer patients achieved a partial response.

To improve the anti-tumor effects of the TLR agonist, we decided to use a combinatorial approach of the TLR4 agonist monophosphoryl lipid A (MPLA) with IFNγ. MPLA is a derivative of LPS; it can be chemically synthesized and displays greatly reduced toxicity while maintaining most of the immunostimulatory activity of LPS. The reduced toxicity of MPLA is attributed to the altered signaling downstream of TLR4 activation compared to LPS, resulting in decreased induction of inflammatory cytokines (*26*). MPLA is used as an adjuvant in commercially available vaccines against cervical cancer and shingles (*27, 28*). IFNγ is a cytokine essential for innate and adaptive immunity. It can promote macrophage activation, enhance antigen presentation, coordinate lymphocyte-endothelium interaction, regulate Th1/Th2 balance, and control cellular proliferation and apoptosis (*29, 30*). Furthermore, IFNγ 1b (a variant of IFNγ) is approved by the FDA to treat chronic granulomatous disease and osteopetrosis (*31, 32*). Here, we describe how injection of MPLA with IFNγ provided a collaborative stimulus that reprogrammed TAMs to become tumoricidal and activated cytotoxic T cells to elicit not only local tumor control but also a systemic immune response to control metastatic breast cancer and ovarian cancer in mice.

## Results

### MPLA with IFNγ reprograms tumor-associated macrophages to tumoricidal macrophages

To explore strategies to induce the tumoricidal activity of macrophages, we first evaluated whether TAMs, which are already polarized to a tumor-promoting phenotype, were plastic enough to be reprogrammed to become tumoricidal. TAMs and tumor cells were isolated from the primary tumors of MMTV-PyMT mice (a mouse model of luminal B breast cancer) (Fig. S1A, S1B), co-cultured and treated with a TLR agonist (LPS or two TLR agonists that have been used clinically: MPLA or polyI:C) alone or in combination with IFNγ. None of the TLR agonists (alone or combined with IFNγ) affected PyMT cancer cell viability in the absence of macrophages (Fig. S1C). However, TAMs activated with IFNγ together with either LPS or MPLA—both TLR4 agonists—killed ~90% of cancer cells in 48 hours in culture (p<0.001, Fig. 1A, B, Movies S1, S2). In contrast, treating macrophages with polyI:C (a TLR3 agonist) combined with IFNγ, or treatment with LPS, MPLA, polyI:C, or IFNγ alone did not induce the tumoricidal activities of the macrophages (Fig. 1A, B).

**Fig. 1.**
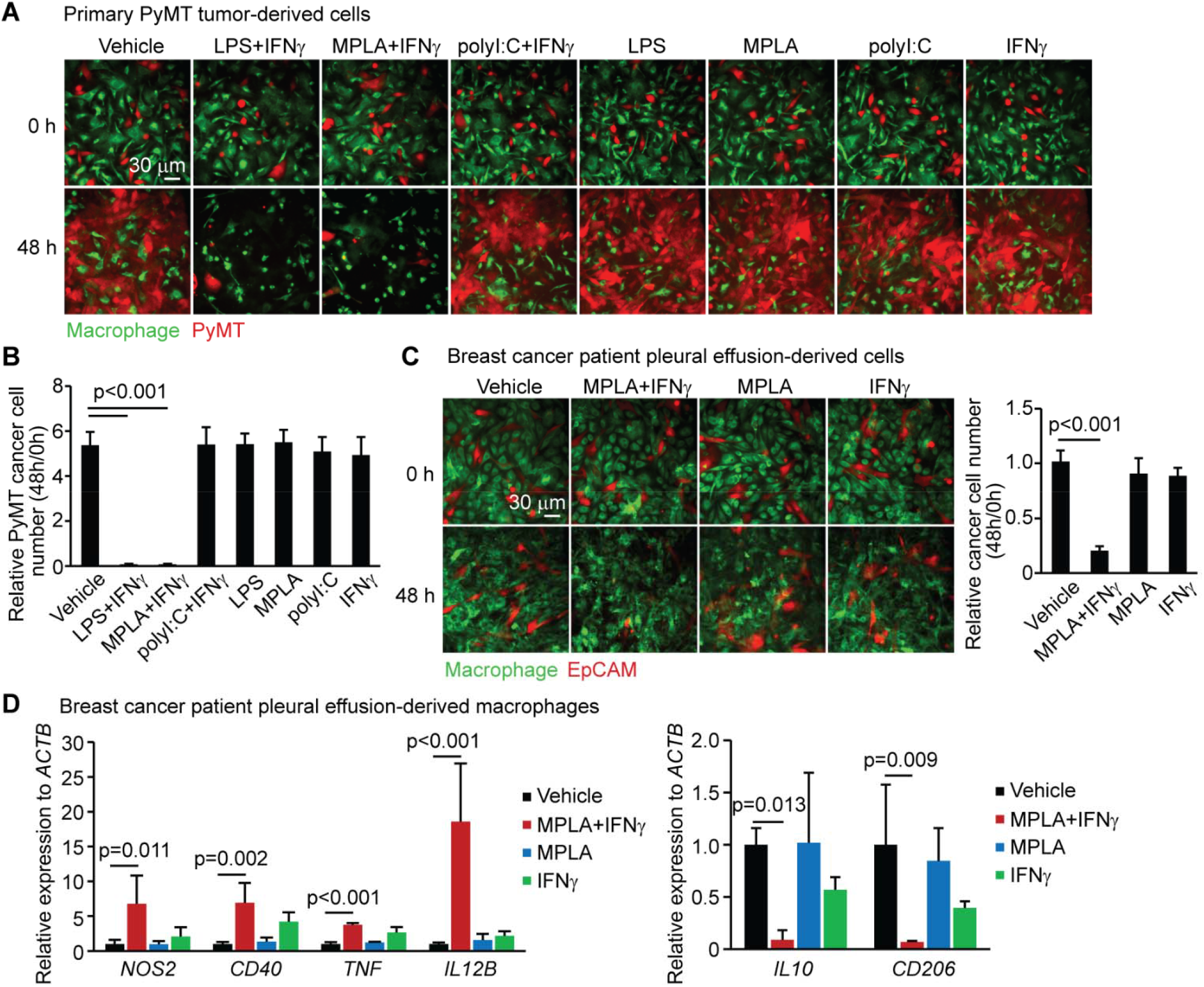
MPLA with IFNγ induces tumoricidal effects of breast tumor-associated macrophages (TAMs) from mice and patients. (**A**) Macrophages and PyMT tumor cells were isolated from the primary tumors of MMTV-PyMT (BL/6) mice and co-cultured for 48 hours in the presence or absence of the indicated treatments. Graph depicts the average results of three independent experiments. (**B**) Statistic analysis of relative PyMT cancer cell numbers for panel A. (**C**) Macrophages and EpCAM^+^ cancer cells were isolated from pleural effusions of breast cancer patients and co-cultured under indicated conditions. EpCAM^+^ cancer cells did not proliferate during co-culture. Graph depicts the average results from three patients. For both A and C, macrophages were stained with CellTracker™ Deep Red Dye, while tumor cells were stained with CellTracker™ CMFDA Dye. Deep Red dye is falsely colored green for clarity. LPS (100 ng/ml), MPLA (100 ng/ml, low-toxicity derivative of LPS; the form we used was produced by chemical synthesis and contains 6 fatty acyl groups), polyI:C (1 μg/ml), mouse IFNγ (33 ng/ml), or human IFNγ (50 ng/ml) was added at 0 hour. (**D**) Expression of genes in pleural effusion-derived macrophages was determined by RT-qPCR 24 hours after indicated treatments were initiated. Graph depicts the average results from three patients. *NOS2*, *CD40*, *TNF*, and *IL12B* (p40 subunit of IL-12) are markers of tumoricidal macrophages. *IL10* and *CD206* are markers of TAMs. The relative expression to *ACTB* was normalized to vehicle, which was set to 1. One-way ANOVA was performed for all panels and mean with standard deviation error bars (mean±s.d.) was presented for each experimental group.

Previously, it had been reported that iNOS is required for the tumoricidal activities of macrophages (*33*). Consistently, only the two combination treatments causing tumoricidal TAM activity—IFNγ combined with either LPS or MPLA—induced iNOS expression (Fig. S1D). Furthermore, inhibition of iNOS activity with L-NIL blocked the tumoricidal effect of macrophages activated by MPLA combined with IFNγ (Fig. S1E). Thus, TAMs can be reprogrammed to become tumoricidal using IFNγ combined with either LPS or MPLA, and iNOS activity is required for their tumoricidal effects.

Recently, CD163^+^ TAMs from the pleural effusions of lung cancer patients were shown to upregulate iNOS expression upon *in vitro* treatment with pseudomonas aeruginosa-mannose-sensitive hemagglutinin (*34*), suggesting that TAMs in patients can be reprogrammed. To formally test this, we isolated macrophages and cancer cells from the metastatic pleural effusions of breast cancer patients (Fig. S1F–H) to determine whether human TAMs could be activated by MPLA with IFNγ to kill patient-derived breast cancer cells. Importantly, MPLA with IFNγ was indeed able to reprogram the pleural effusion-derived macrophages, resulting in the killing of 80–90% of the breast cancer cells isolated from the corresponding patient *in vitro* (p<0.001, Fig. 1C). To characterize the markers associated with the *in vitro* reprogramming of TAMs to tumoricidal macrophages, we next used RT-qPCR. We found that MPLA combined with IFNγ increased the expression of tumoricidal macrophage markers (*NOS2* [the gene coding for iNOS], *CD40*, *TNF,* and *IL12B*) and reduced the expression of TAM markers (*IL10* and *CD206*) in the patient-derived macrophages (Fig. 1D). The upregulation of *Tnf* and *Il12b* and the downregulation of *Il10* and CD206 were also shown in primary PyMT tumor-derived macrophages after MPLA+IFNγ treatment (Fig. S1I, S1J). In aggregate, our data show that the TLR4 agonist MPLA combined with IFNγ reprogrammed mouse and human TAMs to tumoricidal macrophages *in vitro*.

### MPLA with IFNγ suppresses breast tumor growth and lung metastasis

We next sought to determine whether TLR4 agonists combined with IFNγ could control cancer cells *in vivo*, in the context of a complex tumor microenvironment. LPS has severe side effects when given *in vivo* (*35*), but MPLA has been used as adjuvant in vaccines against cervical cancer and shingles (*27, 28*). We therefore tested the effects of intratumoral injection of MPLA with IFNγ into transplanted MMTV-PyMT (“PyMT” for short, luminal B subtype) or 4T1 (triple negative subtype) breast tumors: two models of metastatic breast cancer. We applied 1 μg MPLA with different doses of IFNγ to each tumor and compared their inhibitory effects on tumor growth to optimize the doses. The growth of PyMT tumors treated with MPLA + 0.33, 1, or 3 μg IFNγ was almost equally suppressed compared to controls (Fig. S2A), and the growth of 4T1 tumors treated with MPLA + 3 or 9 μg IFNγ was inhibited similarly (Fig. S2B). Thus, we utilized 1 μg MPLA with 1 μg IFNγ for the intratumoral injection of PyMT tumors, and 1 μg MPLA with 3 μg IFNγ for the intratumoral injection of 4T1 tumors.

This combined MPLA+IFNγ treatment suppressed primary tumor growth in both models (Fig. 2A, 2C). MPLA is a derivative of LPS, which we and others have shown can promote metastasis via activation of neutrophils (*36, 37*), including through the induction of metastasis-promoting neutrophil extracellular traps (NETs) (*38, 39*). However, the treatment did not enhance the levels of NETs in the plasma of PyMT or 4T1 tumor-bearing mice (Fig. S2C, S2D). Importantly, MPLA with IFNγ also did not enhance metastasis: rather, the treatment significantly inhibited lung metastasis. Microscopy of lung sections showed that all vehicle-treated mice had detectable metastasis (Fig. 2B, 2D). In contrast, only two out of five PyMT tumor-bearing mice (Fig. 2B) or four out of nine 4T1 tumor-bearing mice (Fig. 2D) had detectable metastases with the MPLA+IFNγ treatment. We therefore speculate that the dose of MPLA used to activate macrophages is too low to lead to the excessive NET formation that can drive metastasis. Importantly, the suppressive effect of MPLA with IFNγ was collaborative: IFNγ alone inhibited breast tumor growth and lung metastasis, but less than the combination treatment, and MPLA alone had no effect. The ability to reduce metastasis suggests that local injection into the tumor microenvironment can trigger a systemic anti-tumor response. Since intratumoral injection is not the standard of care for any cancer type and since not all tumors have a suitable injectable site, we next tested whether systemic administration of MPLA with IFNγ could control tumor growth. We found that intraperitoneal injection of MPLA (30 μg) with IFNγ (30 μg), similar to intratumoral injection, also reduced breast PyMT tumor growth (Fig. S2E, S2F), and importantly, there were no obvious toxicities.

**Fig. 2.**
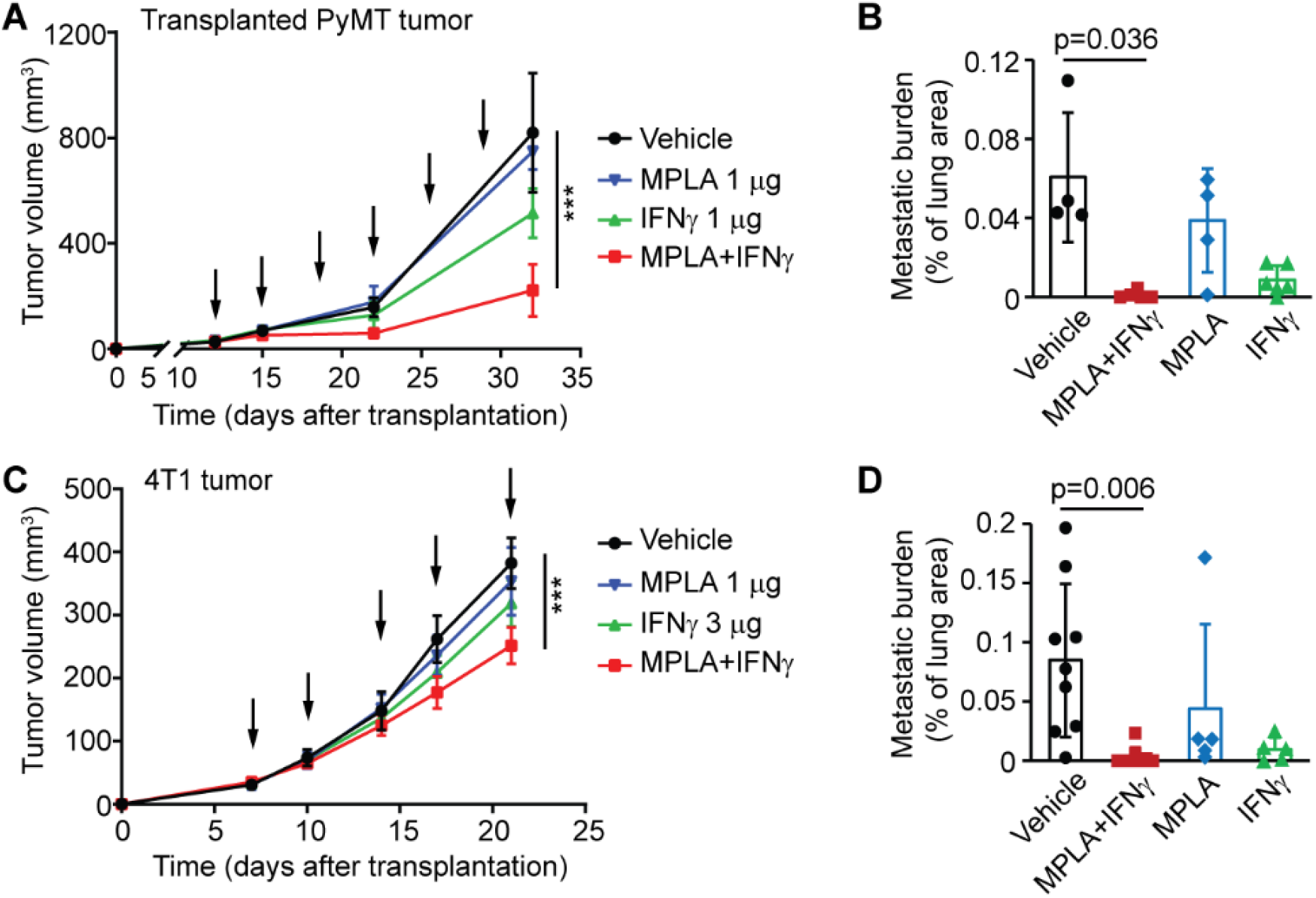
Intratumoral administration of MPLA with IFNγ suppresses breast tumor growth and lung metastasis in mice. (**A**) Tumor growth curve of transplanted PyMT tumors. A total of 2×10^5^ cancer cells isolated from the primary tumors of MMTV-PyMT mice were orthotopically injected into the 4^th^ mammary fat pad of BL/6 mice to obtain PyMT tumors. Arrows indicate time of treatments (twice per week); 1 μg MPLA, 1 μg mouse IFNγ, or the two together were used per tumor. N=8 tumors per group. (**B**) Lung metastatic burden of PyMT tumor-bearing mice was determined by H&E staining. Lung tissues were collected three days after last treatment. N=4–6 mice per group. (**C**) Tumor growth curve for 4T1 tumors: 5×10^4^ 4T1 tumor cells were transplanted into the 4^th^ mammary fat pad of BALB/c mice to obtain 4T1 tumors. Arrows indicate time of treatments (twice per week); 1 μg MPLA, 3 μg mouse IFNγ, or the two together were used per tumor. N=8 tumors per group. (**D**) Lung metastatic burden of 4T1 tumor-bearing mice as determined by H&E staining. Lung tissues were collected three days after last treatment. N=5–9 mice per group. Graphs are representative of at least two independent experiments. One-way ANOVA was performed for the tumor volume analysis at the end time point and for the lung metastasis analysis (mean±s.d.), and because the s.d. was significantly different between the groups (unequal group variances), Welch’s ANOVA was used for Fig. 2A, 2B, and 2D. *** p<0.001, compared to vehicle.

### MPLA with IFNγ stimulates chemokine secretion and type I interferon signaling

To explore how the combination of MPLA with IFNγ reprograms the tumor microenvironment to control tumor growth and metastasis, we focused on the changes induced during the early phases of treatment. We performed cytokine arrays on treated PyMT tumors by collecting tumors 24 hours after the second intratumoral injection (Fig. 3A). To measure cytokines, tumors were digested with collagenase/hyaluronidase and centrifuged, and the cytokines released to the supernatant were collected. Using this method, we found that MPLA combined with IFNγ induced the secretion of a large number of chemokines responsible for immune cell recruitment, including CCL5 (which recruits leukocytes, including monocytes, to inflammatory sites), CXCL9 (which recruits lymphocytes, particularly T cells), CXCL13 (a B lymphocyte chemoattractant), CCL12 (also known as monocyte chemotactic protein 5), and CXCL5 (which recruits neutrophils) (Fig. 3B). In addition, we detected increased CD40 (a co-stimulatory protein on antigen-presenting cells), IL-12 (which enhances the cytotoxic activity of NK cells and CD8^+^ T lymphocytes), and BAFF (B-cell activating factor) (Fig. 3B). We further confirmed by enzyme-linked immunosorbent assay (ELISA) the increased secretion of CXCL9 and CCL5 after combined treatment with MPLA+IFNγ (Fig. S3A, S3B). Together, these data indicate that the immune microenvironment was dramatically altered by the combined treatment of MPLA with IFNγ in breast PyMT tumors.

**Fig. 3.**
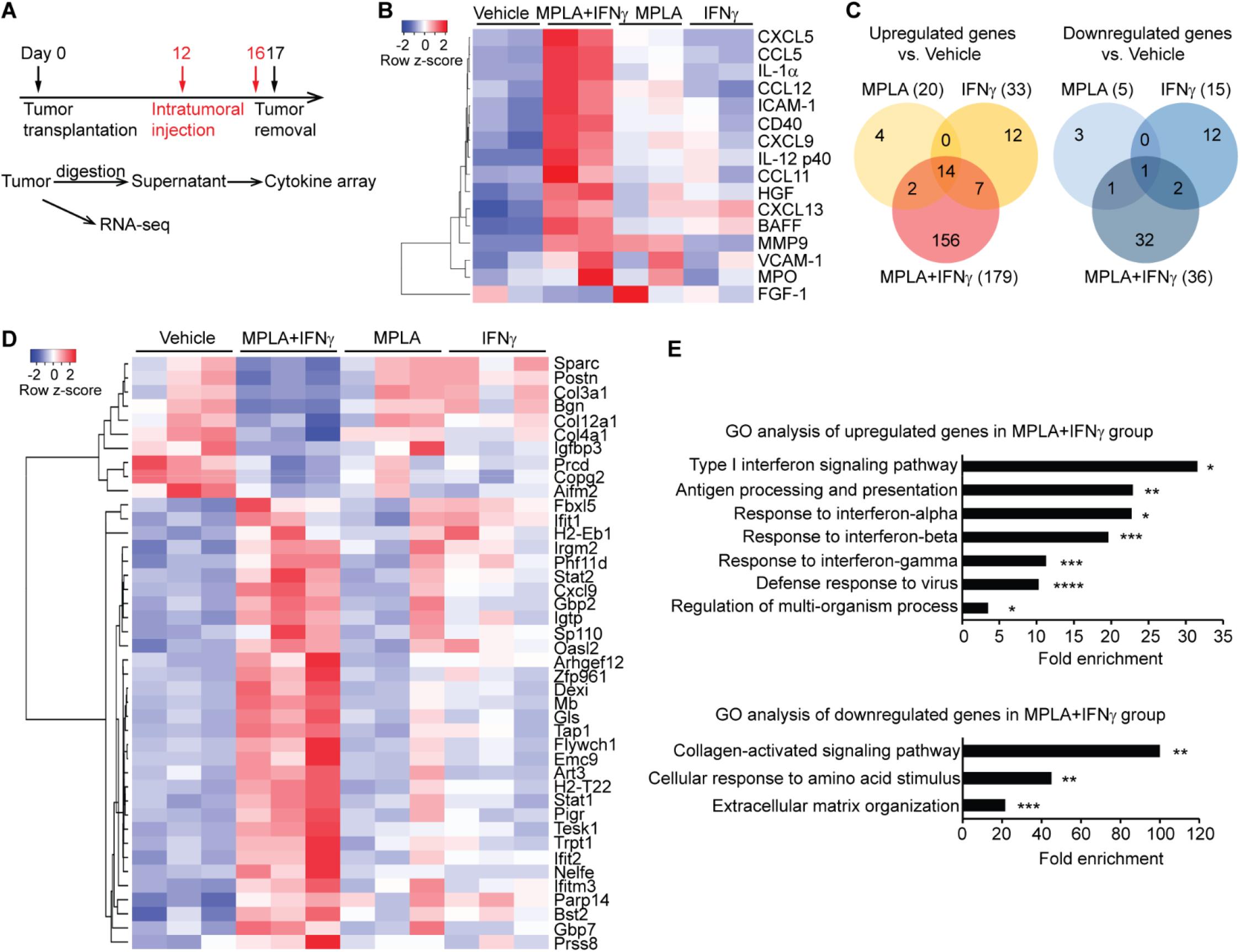
MPLA with IFNγ enhances secretion of chemokines and stimulates the type I interferon signaling pathway in PyMT tumors. (**A**) Experimental design for obtaining cytokine array and RNA-seq data; 1 μg MPLA, 1 μg mouse IFNγ, or the two together were used per PyMT tumor. The treated tumors were collected 24 hours after the second intratumoral injection. (**B**) Heatmap of the secreted cytokines that were changed by the combination treatment (MPLA+INFγ) compared to vehicle. (**C**) Venn diagrams of upregulated and downregulated genes in tumors after the indicated treatments. (**D**) Heatmap of the top genes that were uniquely changed in the MPLA+INFγ treatment group compared to the vehicle group. (**E**) Gene ontology analysis of the genes that were up- or down-regulated in the MPLA+INFγ group. *FDR<0.05, **FDR<0.01, ***FDR<0.001, ****FDR<0.0001, compared to vehicle.

We next explored the effect of combined MPLA+IFNγ on gene regulation by analyzing RNA-seq data (Fig. 3C). There were only 20 upregulated genes and 5 downregulated genes after treatment with MPLA alone, 33 upregulated genes and 15 downregulated genes after treatment with IFNγ alone, but 179 upregulated genes and 36 downregulated genes after the combined MPLA+IFNγ treatment (Data file S1). Thus, the combined treatment was clearly collaborative. Expression of IFNγ-inducible genes was promoted by IFNγ alone and by MPLA+IFNγ (Data file S1). In addition, the combination of MPLA with IFNγ enhanced the expression of type I interferon (interferon-α/β)-inducible genes, including *Stat1*, *Stat2*, *Irgm2*, *Gbp2*, and *Cxcl9* (Fig. 3D). These genes promote oxidative killing, antimicrobial response, and effector T cell response and stimulate the transcription of MHC class II genes, which are critical for the presentation of extracellular pathogens (*40–42*). MPLA with IFNγ also uniquely decreased the expression of extracellular matrix (ECM)-associated genes (*Col3a1*, *Col4a1*, *Col12a1*, and *Igfbp3*) (Fig. 3D). These genes promote cancer cell growth and migration and can suppress the proliferation and cytotoxic activity of T cells (*43, 44*). The altered transcription of selected type I interferon-inducible and ECM-associated genes was verified in PyMT tumors by RT-qPCR and was also shown to occur in combination-treated 4T1 tumors by RT-qPCR (Fig. S3C–F). These results indicate that MPLA with IFNγ activated immune cells and also had wider effects on the tumor microenvironment.

To investigate the functional implications of the altered gene programs, we performed a gene ontology (GO) analysis (Fig. 3E). This analysis revealed enhanced immunostimulatory responses, such as type I interferon signaling, antigen processing and presentation, and defense response to virus, and showed inhibited immunosuppressive responses, such as collagen-activated signaling and ECM organization. Overall, the cytokine array data and RNA-seq data demonstrated that MPLA with IFNγ promoted the expression of genes involved in effector immune cell recruitment and stimulated the type I interferon signaling pathway. These results show that the combination treatment is necessary to induce the changes in the cytokine environment and gene expression that leads to strong anti-tumor responses.

### MPLA with IFNγ reprograms tumor-associated macrophages and activates cytotoxic T cells

Since the GO analysis revealed that pathways involved in immune responses were specifically upregulated after MPLA+IFNγ treatment, we next wanted to elucidate the contributions of immune cells to the anti-tumor effects. By flow cytometry (Fig. S4), we found that as a percentage, there were almost twice as many immune (CD45^+^) cells had infiltrated into PyMT tumors after intratumoral injection of MPLA+IFNγ compared to vehicle-treated tumors (Fig. S5A). Among the immune cells (S5B-J), the proportion of B cells was decreased (S5F), whereas the proportions of CD4^+^ T cells (Fig. S5D) and inflammatory monocytes (Ly6C^+^Ly6G^−^, p=0.002, Fig. 4A) were increased. The infiltration of macrophages in PyMT tumors (F4/80^+^, p<0.001, Fig. 4B) was enhanced. Not only did the tumors contain more macrophages, but the proportion of macrophages that expressed iNOS (p<0.001, Fig. 4C) or CD40 (p<0.001, Fig. 4D) was increased, while the proportion of macrophages that expressed CD206 was reduced (from 50% to 16%, p<0.001, Fig. 4E), as measured by flow cytometry. Consistently, the numbers of iNOS^+^ or CD40^+^ cells were also increased by MPLA+IFNγ, as determined by immunofluorescence (IF) staining of tumor sections (Fig. 4F, 4G). Together, these results suggest that MPLA with IFNγ had reprogrammed TAMs to tumoricidal macrophages in the mouse breast tumors.

**Fig. 4.**
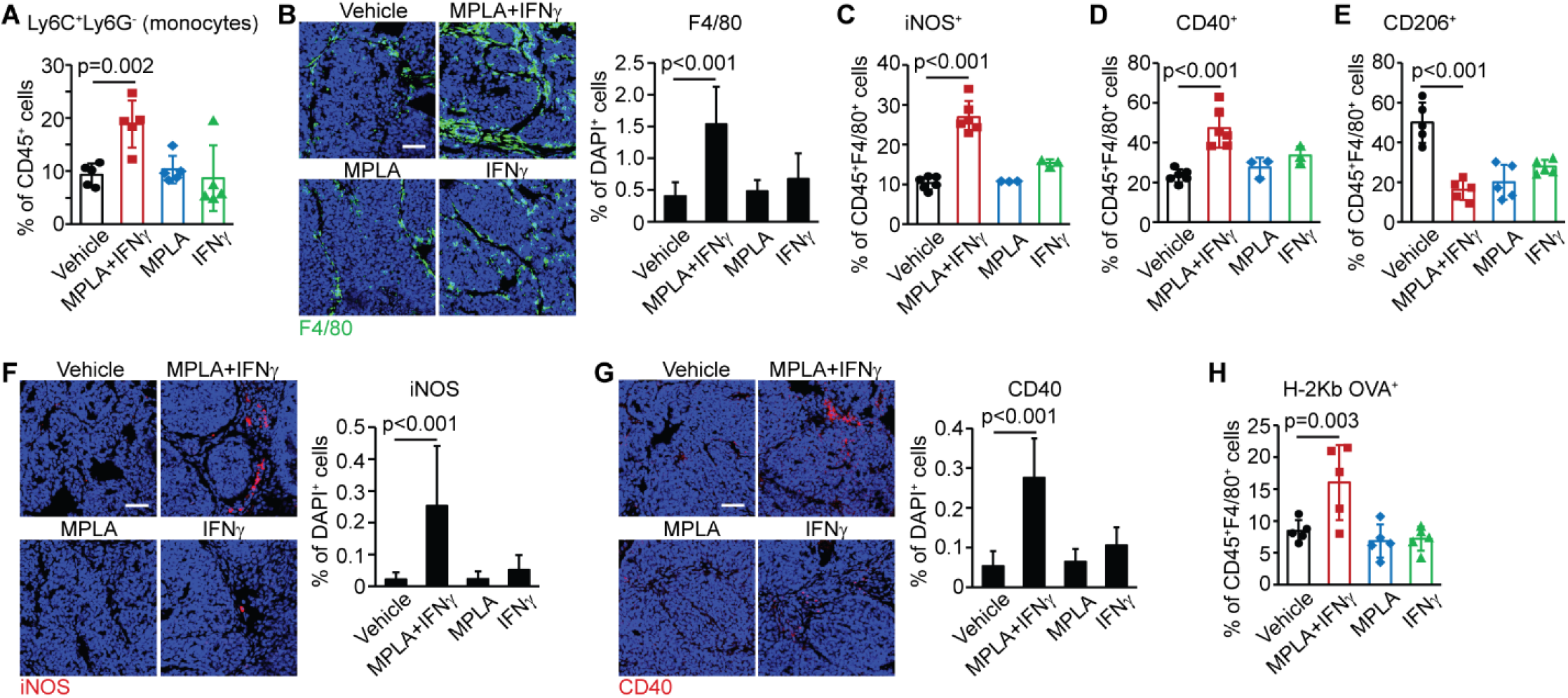
MPLA with IFNγ reprograms TAMs to be tumoricidal in breast tumors. (**A**) Flow cytometry of inflammatory monocytes in PyMT tumors after the last (6^th^) intratumoral injection of the indicated agents. N=5 mice per group. (**B**) IF staining of macrophages (F4/80^+^) in PyMT tumors. In total, 6–14 field of views (FOVs) were evaluated from four mice per group. Scale bar: 50 μm. (**C–E**) The percentage of tumoricidal macrophages (iNOS^+^ or CD40^+^) and TAMs (CD206^+^) of total macrophages were identified by flow cytometry. N=3–6 mice per group. (**F, G**) IF staining of iNOS^+^ (**F**) and CD40^+^ cells (**G**) in breast PyMT tumors. In total, 6–14 FOVs were evaluated from four mice per group. Scale bar: 50 μm. (**H**) Antigen-presenting activities of macrophages were determined by the percentage of H2-Kb OVA^+^ cells in PyMT-chOVA tumors: 2×10^5^ cancer cells isolated from primary tumors of MMTV-PyMT-chOVA mice were transplanted into BL/6 mice to obtain tumors. N=5 mice per group. For all figure panels, the analysis was performed two days after the last (6^th^) intratumoral injection. Graphs are representative of at least two independent experiments. One-way ANOVA was performed for all panels (mean±s.d.), and due to unequal group variances, Welch’s ANOVA was used for Fig. 4B, 4F and 4G.

We next used a variant of the MMTV-PyMT model, the MMTV-PyMT-chOVA model (*45, 46*), to investigate the effect of the combined treatment on MHC class I antigen-presenting activities. Cancer cells in the MMTV-PyMT-chOVA model express the ovalbumin (OVA) peptide SIINFEKL as a model tumor antigen, allowing measurement of MHC class I presentation of OVA. In the tumors of these mice, there was an almost two-fold increase in the percentage of H-2Kb OVA^+^ macrophages (p=0.003, Fig. 4H), H-2Kb OVA^+^ dendritic cells (Fig. S5K), and CD103^+^CD11c^High^CD11b^High^ cells (Fig. S4, S5L) after MPLA+IFNγ treatment. Consistent with the increased antigen presentation, the MPLA+IFNγ treatment significantly increased the infiltration of cytotoxic T lymphocytes (CD8a^+^, p<0.001, Fig. 5A) and the abundance of activated cytotoxic T cells (CD107a^+^ or PD1^+^) in tumors (Fig. 5B, 5C). Importantly, the combined treatment also promoted the production of effector memory T cells (CD44^+^CD62L^−^) (p=0.029, Fig. 5D) and enhanced T cell aggregation in the lungs of tumor-bearing mice (Fig. S5M). Since effector memory cells persist and can provide future protective immunity, together, these data provide evidence that MPLA with IFNγ has the ability to reverse immune suppression in the tumor microenvironment and suggest that the treatment can lead to the activation of lasting adaptive T cell immune responses to protect against future metastases.

**Fig. 5.**
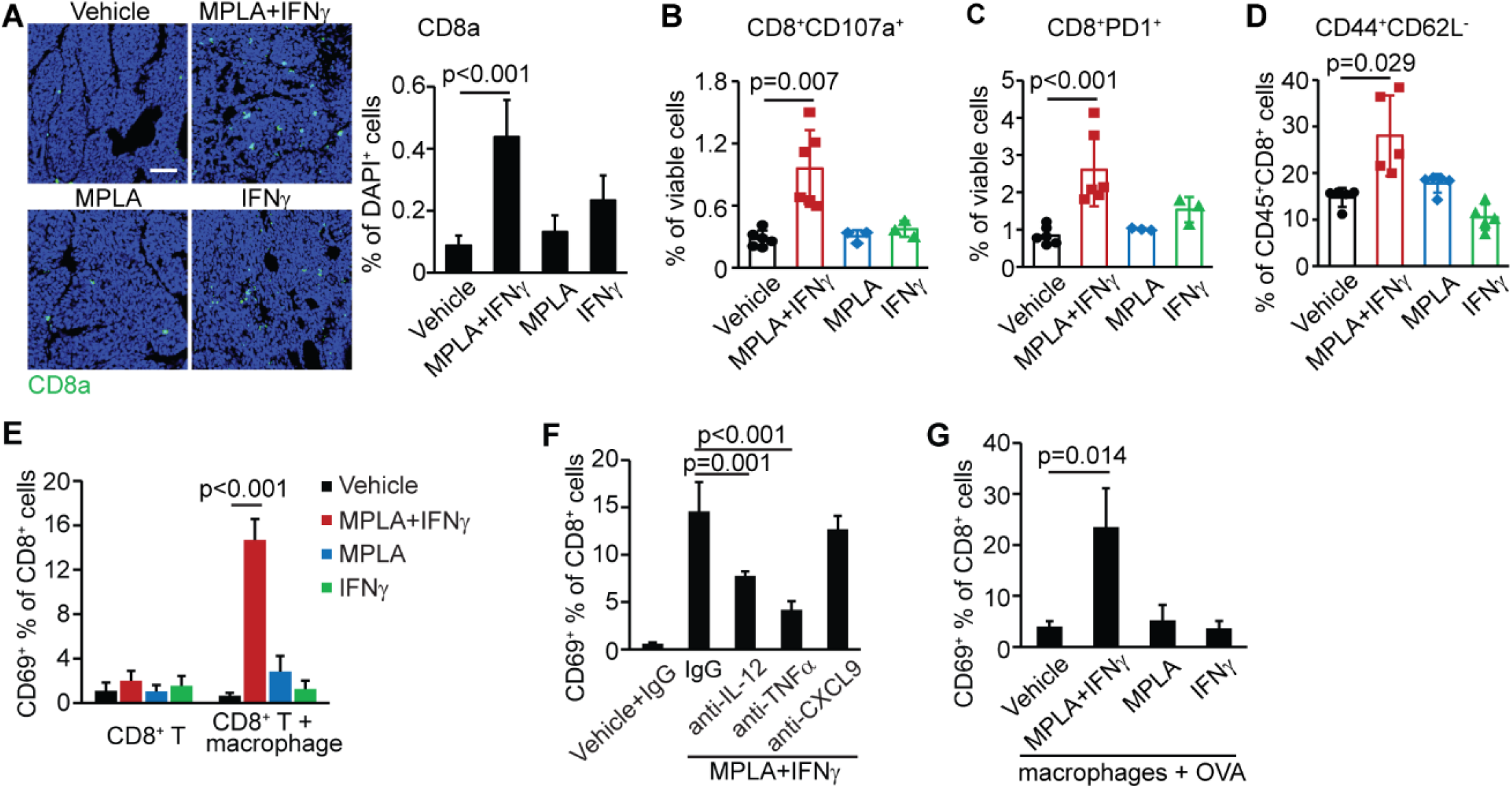
MPLA with IFNγ can activate cytotoxic T cells through macrophage-secreted IL-12 and TNFα. (**A**) IF staining of CD8^+^ T cells in breast PyMT tumors. In total, 5–9 FOVs were evaluated from four mice per group. Scale bar: 50 μm. (**B–D**) The percentages of cytotoxic T cells (CD8^+^CD107a^+^, **B**), PD1^+^CD8^+^ T cells (**C**), and effector memory CD8^+^ T cells (CD44^+^CD62L^−^, **D**) in breast PyMT tumors were identified by flow cytometry. For panels A–D, the analysis was performed two days after the last (6^th^) intratumoral injection. (**E–G**) CD69 expression of CD8^+^ T cells in *in vitro* co-culture systems was identified by flow cytometry. (**E**) CD8^+^ T cells isolated from the spleens of PyMT tumor-bearing mice were cultured with or without macrophages derived from the bone marrow of the same mice, and MPLA (100 ng/ml) (or mouse IFNγ [33 ng/ml] or a combination of the two) was added to the cell cultures immediately after the seeding of T cells. (**F**) CD8^+^ T cells were cultured with macrophages similarly to (E), and the neutralized antibodies of IL-12, TNFα, or CXCL9 were added to co-cultures 30 minutes before adding MPLA with IFNγ. (**G**) MPLA with IFNγ promotes the antigen-presenting activity of macrophages to CD8^+^ T cells. Macrophages derived from the bone marrow of PyMT tumor-bearing mice were activated with MPLA (or IFNγ or a combination of the two) for 16 hours, then incubated with OVA-Q4H7 peptide (10^−9^ M) for 2 hours. The pretreated macrophages were washed with PBS and added to the CD8^+^ T cells that had been isolated from the spleens of OT1 mice. One-way ANOVA was performed for all panels (mean±s.d.), and due to unequal group variances, Welch’s ANONA was used for Fig. 5A, 5B, 5D, and 5G.

### MPLA with IFNγ can activate cytotoxic T cells through macrophage-secreted cytokines

Our *in vivo* data strongly suggested that MPLA with IFNγ treatment activated cytotoxic T cells either directly or indirectly via macrophages. To test these possibilities, we incubated CD8^+^ T cells that were isolated from the spleens of PyMT tumor-bearing mice with MPLA+IFNγ *in vitro*, but found that the combined treatment did not directly stimulate T cell activation (Fig. 5E). In contrast, when macrophages derived from the bone marrow of the same mice and T cells were co-cultured, MPLA with IFNγ increased the expression of CD69 on CD8^+^ T cells (p<0.001, Fig. 5E). Together, these results suggest that MPLA with IFNγ can stimulate cytotoxic T cell functions via macrophages. Since we had found that MPLA with IFNγ promoted the mRNA expression of two T cell activators, *IL12B* and *TNF*, in breast cancer patient pleural effusion-derived macrophages (Fig. 1D) and primary PyMT tumor-derived macrophages (Fig. S1I) and also enhanced the secretion of IL-12 and CXCL9 in PyMT tumors (Figs. 3B, S3A), we explored the possible functions of these three cytokines in the communication between macrophages and T cells *in vitro* by treating with neutralizing antibodies (Fig. 5F). The enhanced expression of CD69 was reduced to 54% by treatment with the IL-12 blocking antibody (p=0.001) and to 39% by the TNFα blocking antibody (p<0.001), indicating that MPLA with IFNγ activates cytotoxic T cells through the macrophage-secreted cytokines IL-12 and TNFα. Blocking CXCL9 had no effect on CD69 expression, consistent with a likely role in the recruitment rather than the activation of T cells.

To further determine how MPLA+IFNγ-activated macrophages affect T cells, we next challenged CD8^+^ T cells isolated from OT1 mice with OVA peptide-incubated macrophages. OT1-derived T cells recognize cells with the MHC class I-bound OVA peptide. We found that the OT1 CD8^+^ T cells were activated by the combination of OVA and MPLA+IFNγ-treated macrophages (p=0.014, Fig. 5G), while MPLA or IFNγ alone had no effect. These results show that MPLA with IFNγ can activate macrophages to present antigens to cytotoxic T cells. Taken together, our *in vitro* results show that MPLA combined with IFNγ not only activates macrophages’ tumoricidal effects but also enhances their capacity to activate cytotoxic T lymphocytes.

### Both macrophages and T cells are essential for the anti-tumor effects of MPLA+IFNγ

We next tested which immune cell populations are necessary for MPLA+IFNγ to eliminate metastatic breast cancer. We performed *in vivo* antibody depletion experiments to target either CD4^+^ and CD8^+^cells (through combined treatment with anti-CD4 and anti-CD8 antibodies) or macrophages (through treatment with anti-CSF1R antibodies). Specific cell depletion was confirmed by flow cytometry of immune cells in tumors (Fig. S6A, S6B). In PyMT tumors, T cell depletion reduced the anti-tumor efficacy of MPLA with IFNγ (Fig. S6C, S6D), but we were unable to determine the contribution of macrophages, as anti-CSF1R antibodies was not effective at depleting them (Fig. S6A). In 4T1 tumors, depleting either macrophages or T cells was effective (Fig. S6B), and the depletion of either cell type abolished the suppressive effects of MPLA+IFNγ on both tumor growth and lung metastasis (Fig. 6A, 6B). These results suggested that both the innate and adaptive immune system are required for the anti-tumor environment elicited by MPLA with IFNγ.

**Fig. 6.**
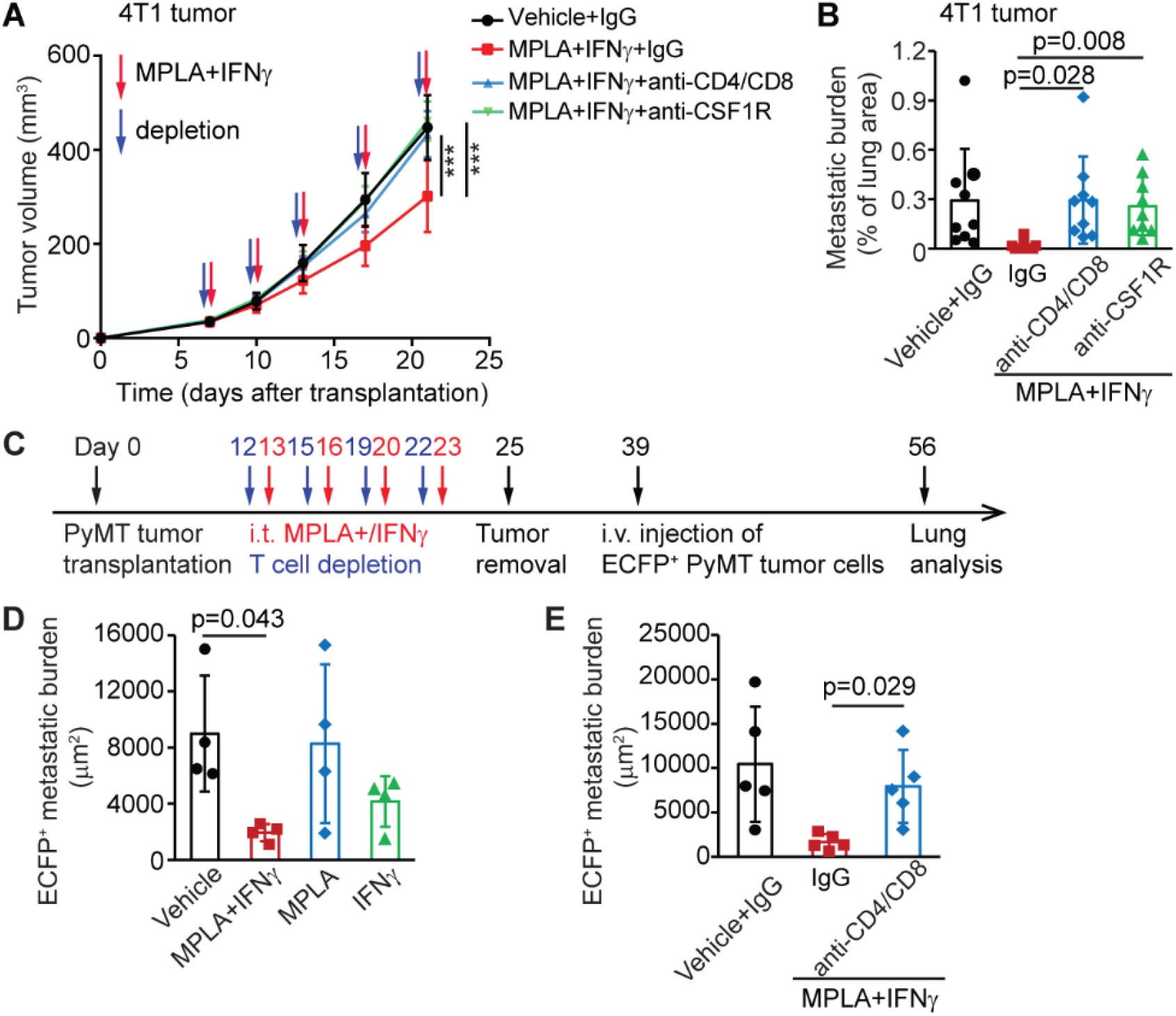
Both the innate and adaptive immune systems are required for the anti-tumor response to MPLA with IFNγ. (**A**) Tumor growth curve of 4T1 tumors after T cell depletion (anti-CD4/CD8) or macrophage depletion (anti-CSF1R). N=8 tumors per group. Immune cell depletion was performed one day before intratumoral injection of MPLA with IFNγ. One-way ANOVA was performed for the tumor volume analysis at the end time point. ***p<0.001 compared to MPLA+IFNγ+IgG. (**B**) Lung metastatic burden of 4T1 tumor-bearing mice was determined by H&E staining. Lung tissues were collected three days after last MPLA+IFNγ treatment. N=9 mice per group. (**C**) Procedure of tumor re-challenge assay with or without T cell depletion. i.t.: intratumoral administration; i.v.: intravenous. (**D–E**) Lung metastasis formed by newly injected breast cancer cells (ECFP^+^) without (**D**) or with (**E**) T cell depletion was determined by IF staining of lung sections for ECFP. ECFP^+^ cancer cells were isolated from the primary tumors of MMTV-PyMT; ACTB-ECFP mice. N=4–5 mice per group. One-way ANOVA was performed for all lung metastasis analyses (mean±s.d.), and due to unequal group variances, Welch’s ANONA was used for Fig. 6B, 6D, and 6E.

Our data suggested that a memory T cell response is activated by MPLA+IFNγ treatment (Fig. 5D). To test whether a long-lasting immune response was indeed induced, tumor-bearing mice had their tumors surgically removed, recovered for two weeks, and were then re-challenged with cancer cells from MMTV-PyMT; ACTB-ECFP mice (Fig. 6C), allowing us to identify the new cells by their expression of enhanced cyan fluorescent protein (ECFP). We found that mice treated with MPLA+IFNγ had a significantly reduced ECFP^+^ metastatic burden, whereas MPLA- or IFNγ-treated mice had no reduction (Fig. 6D). When T cells were depleted during MPLA+IFNγ treatment (Fig. 6C), the inhibitory impact of MPLA with IFNγ on metastasis from the new cancer cells was abolished (p=0.029, Fig. 6E). These data, together with the enhanced generation of functional memory T cells by MPLA with IFNγ (Fig. 5D), indicate that the combination treatment not only reverses immune suppression in the tumors but also triggers anti-tumor memory, which may have a long-lasting effect on metastasis.

### MPLA with IFNγ suppresses metastatic ovarian cancer

Intraperitoneal MPLA+IFNγ treatment was successful in our breast cancer model (Fig. S2F). This finding motivated us to explore this combination treatment for ovarian cancer, for which intraperitoneal chemotherapeutic treatments are used and for which there is a lack of effective treatments for late-stage disease. Furthermore, TAMs contribute to immunosuppressive microenvironment in ovarian cancer (*47, 48*). To determine whether the therapeutic benefit of MPLA with IFNγ might be extended to ovarian cancer, we intraperitoneally injected mice with ID8-p53^−/−^ ovarian cancer cells (*49*), a model that is relevant because Trp53 mutations are found in 94% of human high-grade serous ovarian cancer (*50*). At 7 weeks after ID8-p53^−/−^ cancer cell transplantation (Fig. 7A), the tumor-bearing mice were visibly sick (Fig. S7A) and had bloody ascites in their abdominal cavity (Fig. 7B). All mice had a higher body weight (Fig. S7B) and many metastases in the peritoneum, pancreas, mesentery, and diaphragm (Table S1). However, when treatment with MPLA+IFNγ was initiated 3 weeks after cancer cell transplantation (Fig. 7A), the mice at the 7-week time point looked healthy (Fig. S7A), and only two out of five mice had metastases in the peritoneum (Table S1). Furthermore, MPLA combined with IFNγ decreased the total cell number in the ascites (to just 10% of that found in control mice, p=0.007, Fig. 7C), increased the percentage of immune cells in the ascites (CD45^+^, Fig. S7C), and reduced the proportion of EpCAM^+^ cancer cells (from 15% to 2%, p=0.020, Fig. 7D), indicating that the treatment suppressed the progression of ovarian cancer.

**Fig. 7.**
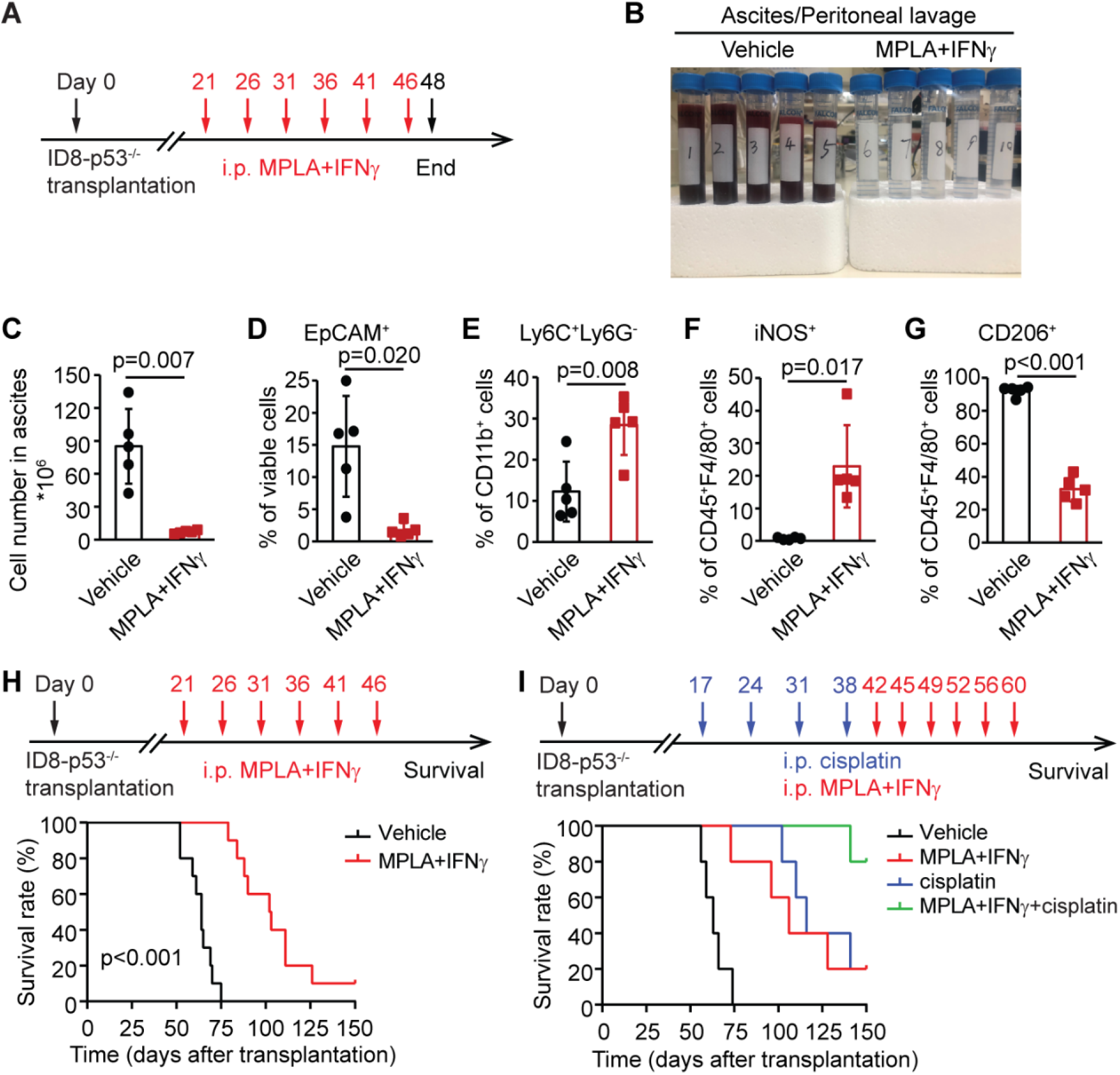
MPLA with IFNγ suppresses metastatic ovarian cancer in mice. (**A**) Experimental design. A total of 5×10^6^ ID8-p53^−/−^ ovarian cancer cells were injected intraperitoneally into C57BL/6 mice. Three weeks later, 10 μg MPLA with 20 μg mouse IFNγ was applied to the mice every five days, for a total of 6 times. Two days after the last treatment (~7 weeks after transplantation), tumor-bearing mice were euthanized and ascites/peritoneal lavage was collected for flow cytometry. i.p.: intraperitoneal administration. (**B**) Metastasis was evident from the appearance of the ascites/peritoneal lavage of the treated mice. (**C, D**) Metastasis was determined by the total cell number (**C**) and percentage of epithelial cancer cells (EpCAM^+^, **D**) in the ascites/peritoneal lavage. (**E–G**) The proportions of monocytes (Ly6C^+^Ly6G^−^, **E**), of tumoricidal macrophages (iNOS^+^, **F**), and of TAMs (CD206^+^, **G**) in ascites/peritoneal lavage were determined by flow cytometry. N=5 mice per group. Graphs are representative of at least two independent experiments. Unpaired t-test was performed (mean±s.d.), and due to unequal group variances, Welch’s t-test was used for Fig. 7C, 7D, and 7F. (**H**) Kaplan-Meier survival curves of ID8-p53^−/−^ tumor-bearing mice treated with MPLA+IFNγ every five days. N=10 mice per group. Log-rank test was used: p<0.0001. (**I**) Kaplan-Meier survival curves of ID8-p53^−/−^ tumor-bearing mice treated with MPLA+IFNγ alone or in combination with cisplatin (10 μg/g body weight) administered as indicated (N=5 mice per group. Log-rank test was used: p<0.0001. Median survival times: Vehicle: 63 days; MPLA+IFNγ: 106 days; cisplatin: 116 days; and cisplatin followed by MPLA+ IFNγ: undefined at 150 days.

MPLA with IFNγ increased the presence of monocytes in the ascites (p=0.008, Fig. 7E), and similar to the breast cancer model, the combination treatment altered the balance between the iNOS^+^ and CD206^+^macrophages (Fig. 7F, 7G): the percentage of iNOS^+^ macrophages increased from 0.7% to 23% (p=0.017), while the percentage of CD206^+^ macrophages decreased from 92% to 33% (p<0.001). Of note, the mouse that had the most metastases in the MPLA+IFNγ group had the lowest percentage of iNOS^+^ macrophages and the highest percentage of CD206^+^ macrophages, supporting a direct link between the ability to reprogram macrophages and reduce metastasis. Strikingly, MPLA+IFNγ treatment prolonged the survival of ID8-p53^−/−^ tumor-bearing mice regardless of whether treatment was started at 3 weeks after tumor inoculation (Fig. 7H), or six weeks after tumor inoculation, when many mice had small amount of bloody ascites (Fig. 7I).

Most patients with ovarian cancer will be treated with chemotherapy, but nevertheless the cancer will often recur (*51*). We therefore compared treatment with MPLA+IFNγ to cisplatin, a widely used chemotherapy for ovarian cancer patients. Cisplatin, given weekly for four weeks starting day 17 after tumor cell inoculation, and MPLA+IFNγ, given twice weekly for three weeks starting day 42, increased the median survival of the mice, largely to the same extent (Fig. 7I). When MPLA+IFNγ was given after four cycles of cisplatin, the median survival was increased compared to the monotherapies (Fig. 7I). Thus, MPLA+IFNγ may offer benefit when given after standard of care chemotherapy. Altogether, our data show that MPLA combined with IFNγ dramatically suppresses metastatic breast and ovarian cancers by reversing the tumor immunosuppressive microenvironment and stimulating the collaborative innate-adaptive anti-tumor immune response.

## Discussion

Breast and ovarian tumors are infiltrated by large numbers of macrophages and T cells. Although these cells can kill cancer cells, they must be activated properly. With this study, we developed a strategy for reprogramming the immune microenvironment of cancer. We revisited the approach used for Coley’s toxin (dead bacteria) and exploited the combined power of the TLR4 agonist MPLA (a derivative of the bacterial component LPS) with the cytokine IFNγ. The combination of MPLA with IFNγ reversed immune suppression in the tumor microenvironment and induced a systemic anti-tumor response. This treatment harnessed the power of macrophages and T cells so that these cells together eliminated metastatic breast and ovarian cancer. This result is in line with previous reports using other approaches to activate macrophages (*18, 19*), strongly supporting the argument for engaging both innate as well as adaptive immunity to improve the efficacy of cancer immunotherapy. Importantly, our strategy for co-stimulating macrophages and T cells did not require knowledge of the tumor antigens.

We found that MPLA with IFNγ stimulated innate immunity, but in addition, the adaptive immune response was critical for the *in vivo* effects of the treatment. We found that MPLA with IFNγ did not stimulate T cells directly *in vitro* and that the activation of CD8^+^ T cells depended on cytokines secreted by macrophages, including IL-12 and TNFα. It is likely that MPLA+IFNγ *in vivo* also induces dendritic cells to produce IL-12 to activate cytotoxic T lymphocyte responses, as such effects have been reported (*52*). Our data are consistent with reports that IL-12 bridges innate and adaptive immunity by enhancing the cytotoxic activity of CD8^+^ T lymphocytes (*53*) and that intratumor adoptive transfer of tumor-specific CD8^+^ T cells engineered with transiently expressed IL-12 mRNA eradicated tumors in melanoma mouse models (*54*). TNFα can promote the activation and proliferation of naïve and effector T cells (*55, 56*). Macrophages from murine tumors, as well as from malignant pleural effusions of breast cancer patients could be reprogrammed to tumoricidal macrophages by MPLA with IFNγ *ex vivo*. Importantly, this suggest that TAMs in the metastatic tumor microenvironment are sufficiently plastic to be reprogrammed to become tumoricidal. Furthermore, the reprogrammed patient-derived macrophages expressed *IL12B* and *TNF*, suggesting that a T cell response might also be activated in patients with cancer.

The collaborative therapeutic effect between MPLA and IFNγ *in vivo* is likely also related to the fact that only the combined treatment induces a type I interferon response in the tumors. Treatment with just IFNγ, which is not a type I interferon, did not induce such a response. Type I interferons play critical roles in anti-cancer immunity (reviewed in (*57–59*)). Important roles include their ability to promote antigen presentation, which is consistent with our data showing increased the antigen-presenting activities of macrophages and dendritic cells after treatment by MPLA with IFNγ, but not by either treatment alone. Type I interferons also skew the adaptive immune system to promote the development of antigen-specific T cell responses and immunological memory. Indeed, we observed that MPLA with IFNγ significantly increased the abundance of activated cytotoxic T lymphocytes and effector memory T cells in tumors. In the tumor re-challenge assay, MPLA+IFNγ-treated mice developed the fewest new metastatic lesions, and T cells were required for this effect, suggesting that the combination treatment triggers long-term anti-tumor T cell memory.

In our experiments*, in situ* injection of 1 μg MPLA (~50 μg/kg body weight) with 1–3 μg IFNγ (~50–150 μg/kg body weight) was sufficient to induce local immune modulation in breast tumors and to invoke a systemic anti-tumor immune response that protected against metastatic disease, either from spontaneously disseminated tumor cells or from cells injected as a re-challenge of the immune system. Potential drawbacks of invoking local immune modulation include the reliance on adequate immune infiltrates available for reprogramming and the requirement for suitable injectable sites in the tumor. As an alternative, we administrated MPLA with IFNγ intraperitoneally. Although this approach required higher doses (10–30 μg, and ~0.5–1.5 mg/kg body weight), these are still low compared to other systemically administered antibodies and cytokines.

MPLA and IFNγ are already FDA-approved biologics: MPLA as an adjuvant is used in vaccines against cervical cancer and shingles, and a variant of IFNγ (IFNγ 1b) is approved to treat chronic granulomatous disease and osteopetrosis. Importantly, we observed no obvious toxicity or induction of metastasis-promoting NETs at the doses that could induce protective immunity in mice, indicating that this combination treatment could be a strategy for eliminating tumors without causing severe side effects. We therefore envision that MPLA with IFNγ may be applied as a stand-alone treatment or perhaps used in conjunction with chemotherapeutic agents to prevent metastatic recurrence. Metastasis is the leading cause of mortality in cancer patients (*60*). If the elimination of metastasis by MPLA+IFNγ can be translated from our preclinical models to patients, then the approach may significantly improve the survival rates of late-stage breast and ovarian cancer patients.

## Materials and Methods

### Study design

The objective of this research was to develop a new therapeutic strategy to reprogram TAMs to tumoricidal macrophages and to reverse the tumor immunosuppressive microenvironment to an immunostimulatory microenvironment. We examined the anti-tumor effect of MPLA with IFNγ by measuring tumor growth and metastasis in mouse models of metastatic luminal (PyMT) and triple negative (4T1) breast cancers, as well as in an ovarian cancer model (ID8-p53^−/−^). Importantly, we also investigated the ability of MPLA with IFNγ to reprogram TAMs from the malignant pleural effusions of breast cancer patients. We studied the mechanism of the therapy by performing a cytokine array, RNA-seq, and flow cytometry to examine the activation of immune cells and by performing neutralization/depletion experiments to explore the requirement for macrophage-secreted cytokines and innate-adaptive immunity. In all *in vitro* experiments, the average results were taken from at least three independent experiments or three patients. In all *in vivo* experiments, the results are representative of at least two independent experiments (the numbers of mice and statistical tests are listed in the figure legends). Mice were excluded from the study when the breast tumors grew into the abdominal cavity and were randomized before the administration of therapy.

### Animals

OT1 mice were purchased from Jackson Laboratory and MMTV-PyMT mice (on C57BL/6 [referred to hereafter as BL/6] background) were bred at Cold Spring Harbor Laboratory (CSHL). To obtain MMTV-PyMT; ACTB-ECFP mice, ACTB-ECFP mice (on BL/6 background, originally obtained from Jackson Laboratory) were intercrossed with MMTV-PyMT mice. MMTV-PyMT-chOVA mice (*46*) were a kind gift from Dr. Mathew Krummel (University of California, San Francisco).

Six-to eight-week-old female BL/6 and BALB/c host mice were purchased from Jackson Laboratory and acclimated to the animal housing facility for one week prior to performing experiments. All animal experiments were conducted in accordance with procedures approved by the Institutional Animal Care and Use Committee at CSHL and the National Institutes of Health (NIH) Guide for the Care and Use of Laboratory Animals.

### Patient samples

Metastatic pleural effusions from three female breast cancer patients were collected during a standard therapeutic procedure used to palliate symptoms. Patient 1 was 34 years old and the tumor was estrogen-receptor (ER) positive (^+^), human epidermal growth factor receptor (HER)-2 negative (^−^); patient 2 was 35 years old and the tumor was ER^+^/HER2^+^; and patient 3 was 53 years old and the tumor was ER^+^/HER2^−^. All research was performed in accordance with the New York University and CSHL Institutional Review Board guidelines and regulations, and written informed consent was obtained from all patients.

### Isolation of macrophages and tumor cells from pleural effusions of breast cancer patients

Pleural effusions were shipped on ice to CSHL and cells were isolated in less than 24 hours after sample collection as follows: the effusions were spun down at 450 × g for 10 minutes to remove the liquid supernatant, and the cell pellet was washed with Dulbecco’s Phosphate-Buffered Saline (DPBS) twice, then resuspended in Ammonium-Chloride-Potassium (ACK) lysis buffer on ice for 5–10 minutes; an additional spin down at 300 × g for 10 minutes was performed to eliminate the red blood cells. Then, CD326 (EpCAM) microbeads (#130-061-101, Miltenyi Biotec) were applied to enrich for EpCAM^+^ and EpCAM^−^ cells, according to the manufacturer’s instructions. Macrophages were obtained from the EpCAM^−^ cells after 2 hours of adherence to the tissue culture dish and were cultured with 50 ng/ml human macrophage colony-stimulating factor (M-CSF, #CYT-308, ProSpec) in RPMI-1640 (Gibco) containing 20% fetal bovine serum (FBS, Sigma), 200 μM L-glutamine (#25030081, Gibco), 100 μM sodium pyruvate (#11360070, Gibco), and 50 μM 2-mercaptoethanol (#31350010, Gibco). EpCAM^+^ cells were cultured with Dulbecco’s Modified Eagle Medium (DMEM)-F12 (Gibco) containing 20% FBS (Sigma), 2% B27 (#17504044, Gibco), 20 ng/ml human EGF (#CYT-217, ProSpec), and 4 μg/ml insulin (#12585014, Gibco).

### Isolation and culturing of macrophages and cancer cells from primary breast tumors of mice

Macrophages and cancer cells for *in vitro* co-culture assays, Western blot analysis, and MTS assays were isolated from the primary tumors of MMTV-PyMT mice (in BL/6 background). The tumors (6–8 mm in diameter) were mechanically dissociated and digested with collagenase/hyaluronidase (#07912, STEMCELL Technologies) containing DNase I (4 U/ml, #4716728001, Roche) at 37 °C for 45 minutes. After pulse centrifugation (450 × g), the supernatant (mostly immune cells) and cell pellets (mostly cancer cells) were collected. CD11b MicroBeads (#130-049-601, Miltenyi Biotec) were used to enrich CD11b^+^ cells from the supernatant, and macrophages were obtained by allowing the CD11b^+^ cells to adhere to the tissue culture dish for 2 hours (non-adherent cells were removed by washing with PBS). Cancer cells were isolated from the tumor cell pellets and dissociated into single tumor cells with TrypLE Express Enzyme (#12605010, Gibco) containing DNase I at 37 °C for 10 minutes.

Mouse macrophages were cultured with RPMI-1640 (Gibco) containing 20% FBS (Sigma), 200 μM L-glutamine (Gibco), 100 μM sodium pyruvate (Gibco), 50 μM 2-mercaptoethanol (Gibco), and 10 ng/ml mouse M-CSF (#CYT-439, ProSpec). Cancer cells were cultured with DMEM + 10% FBS.

### In vitro macrophage and tumor cell co-culture assay

A 24-well glass bottom plate (#P24G-1.5-13-F, MatTek) was coated with fibronectin (1:100, #F1141, Sigma Aldrich). Macrophages were seeded (1.5×10^5^/well) to the plate and 2 hours later, stained with CellTracker™ Deep Red Dye (1 μM, #C34565, Invitrogen). Tumor cells were stained with CellTracker™ Green CMFDA (5-chloromethylfluorescein diacetate) Dye (1 μM, #C7025, Invitrogen) and seeded (5×10^4^/well) to the plate. Two hours later, LPS (100 ng/ml, #L4391, Sigma Aldrich), MPLA (a low toxicity derivative of LPS; 100 ng/ml, #tlrl-mpls, InvivoGen; this product is produced by chemical synthesis and contains 6 fatty acyl groups), polyI:C (1 μg/ml, #tlrl-picw, InvivoGen), mouse IFNγ (33 ng/ml, #485-MI/CF, R&D Systems), or human IFNγ (50 ng/ml, #285-IF/CF, R&D Systems) was added to the medium, alone or in indicated combinations, and live cell imaging was performed with a Yokagawa spinning disk confocal microscope (Solamere Technology Group) and LiveCell Stage Top Incubation System (Pathology Devices). To inhibit iNOS activity, L-NIL (N^6^-(1-Iminoethyl)-L-lysine hydrochloride, 200 μM, #1139, TOCRIS) was added to the medium before MPLA with IFNγ.

### RNA extraction and quantitative real-time PCR

Total RNA from macrophages or primary tumors was extracted using an RNeasy Mini kit (#74104, Qiagen), following the manufacturer’s instructions. Reverse transcription was performed with TaqMan™ Reverse Transcription Reagents (#N8080234, Invitrogen). Quantitative real-time PCR was performed in a 384-well format on a QuantStudio 6-flex Instrument (Applied Biosystems) using PowerUp™ SYBR™ Green Master Mix (#A25780, Applied Biosystems). Relative quantitation was performed with the 2^(−ΔΔCT)^ method using β-actin expression for normalization. The sequences of PCR primers are listed in Table S2.

### In vivo tumor experiments

PyMT cancer cells for orthotopic transplantation were isolated from the primary tumors of MMTV-PyMT mice or MMTV-PyMT-chOVA mice (BL/6 background). The cancer cells were isolated as described above and were injected immediately into the 4^th^ mammary fat pad of BL/6 mice (2×10^5^ cells/fat pad in 20 μl 1:1 PBS/Matrigel [#356231, Corning]) to obtain the transplanted PyMT tumor model. For 4T1 cells, 5×10^4^ cells/fat pad were transplanted into the 4^th^ mammary fat pad of BALB/c mice to obtain the 4T1 tumor model. Tumor size was measured by caliper, and tumor volume was calculated as volume = (length×width^2^)/2. Treatment was initiated when tumors reached 5–6 mm in size and was given twice a week: 1 μg MPLA/tumor (#vac-mpls, InvivoGen) and indicated doses of mouse IFNγ (#485-MI/CF, R&D Systems) were injected into the tumors to optimize the doses by comparing their inhibitory effects on tumor growth. Because the growth of PyMT tumors treated with MPLA+0.33, 1, or 3 μg IFNγ was almost equally suppressed compared to controls (Fig. S2A) and the growth of 4T1 tumors treated with MPLA+3 or 9 μg IFNγ was inhibited similarly (Fig. S2B), we chose the following doses: 1 μg MPLA with 1 μg IFNγ (or 1 μg MPLA alone or 1 μg IFNγ alone) was used for the intratumoral injection of PyMT tumors, and 1 μg MPLA with 3 μg IFNγ (or 1 μg MPLA alone or 3 μg IFNγ alone) was used for the intratumoral injection of 4T1 tumors. For intraperitoneal administration of PyMT tumors, 30 μg MPLA with 30 μg IFNγ per mouse were used. At the indicated time points, mice were euthanized in a CO_2_ chamber before performing a cardiac perfusion with PBS, and tumors and lungs were removed for analysis.

For the tumor re-challenge experiments (shown in Fig. 6C), transplanted PyMT tumors were treated with MPLA+IFNγ (intratumoral injection) four times. Two days after the last treatment, breast tumors were surgically removed by making a small incision at the skin near the site of the tumor. The tumor along with 4^th^ mammary fat pad was excised and the wound was closed using Relfex 7 mm wound clips (#203-1000, CellPoint Scientific). Two weeks after tumor removal, ECFP^+^ PyMT cancer cells were injected into the tail vein of the mice. The ECFP^+^ PyMT cancer cells were isolated from the primary tumors of MMTV-PyMT; ACTB-ECFP mice as described above, and 5×10^5^ cells (in 100 μl PBS) were injected into the tail vein of the mice whose tumors had been surgically removed. Seventeen days later, the lungs were removed for analysis.

Ovarian cancer cells ID8-p53^−/−^ were cultured as previously described (*49*). A total of 5×10^6^ ID8-p53^−/−^ cells (in 200 μl PBS) were injected intraperitoneally into BL/6 mice. At indicated times, 10 μg MPLA with 20 μg mouse IFNγ was administered intraperitoneally to the mice, for six times total. At the end of the experiment (two days after last treatment), the mice were weighed and intraperitoneally injected with PBS, and the ascites/peritoneal lavage was collected (~12 ml collected per mouse) for cell counting and flow cytometry. Ascites/peritoneal lavage was spun down at 300 × g for 10 minutes, and the cell pellet was resuspended in ACK lysis buffer on ice for 5–10 minutes, then spun down at 300 × g for an additional 10 minutes to eliminate the red blood cells. The total cell numbers were determined by a CellDrop cell counter (DeNovix, Inc.). Cisplatin (10 μg/g body weight, #JM-1550-1000, MBL) was administrated intraperitoneally to the mice every week, for a total of four times. MPLA (10 μg) with mouse IFNγ (20 μg) was administered as indicated. Mice were euthanized when the presence of ascites was evident and affected the mobility of the mouse.

### Lung metastasis analysis

Lungs were placed in PBS in a vacuum desiccator for 2 hours, then fixed in 4% paraformaldehyde (PFA) for 4 hours, infiltrated with 30% sucrose in PBS for 48 hours, and embedded in OCT compound. The blocks were then placed in a custom-made cutting chamber with razor blade inserts every 2 mm, and the block was cut into 2 mm sections. These sectioned lung pieces were then re-embedded in fresh OCT and sent to the Histology Core at CSHL for cross-sectional cuts from the entire lung tissue and hematoxylin and eosin (H&E) staining. The metastatic burden was calculated as the percentage of metastasis/lung area, determined using Aperio eSlide Capture Devices software (Leica Biosystems).

### Cytokine array

Twenty-four hours after the second intratumoral injection, transplanted PyMT tumors were collected, weighed, and digested with collagenase/hyaluronidase (25 mg tumor/1 ml enzyme) containing DNase I at 37 °C for 30–45 minutes. The supernatant that contained secreted cytokines was collected and spun down to remove cells and cell debris (12,000 × g, 30 minutes). Supernatant samples (1 ml) and a Proteome Profiler Mouse XL Cytokine Array Kit (#ARY028, R&D Systems) were used according to the manufacturer’s instructions. Films were scanned, and spots were analyzed using ImageJ software. The heatmap was generated with http://www2.heatmapper.ca/expression/.

### RNA sequencing

Twenty-four hours after the second intratumoral injection, transplanted PyMT tumors were collected and total RNA was extracted using an RNeasy Mini Kit (#74104, Qiagen). RNA-seq libraries were prepared using a TruSeq Stranded mRNA Library Prep Kit (#20020594, Illumina) and sequenced using a NextSeq500 High Output Illumina platform with 76 base single end reads at the CSHL Next Generation Sequencing Core Facility, which yielded an average of 33.3 million filtered reads per sample. FASTQ sequencing files were processed and analyzed using the Galaxy server hosted by the CSHL Bioinformatics Core Facility. Reads were aligned to the GRCm38/mm10 reference mouse genome. Gene read counts were determined using featureCounts (*61*). Differential gene expression analysis was performed using DESeq2, with a 0.1 false discovery rate (FDR)-adjusted p-value cutoff for significance. DESeq2-normalized read counts of the top genes that were uniquely changed in the MPLA+IFNγ group were visualized as a heatmap using http://www2.heatmapper.ca/expression/. Genes that were uniquely upregulated or downregulated in the MPLA+IFNγ group were used as the input for biological process gene ontology analysis (http://geneontology.org/).

### Flow cytometry

Two days after the last (6^th^) intratumoral injection, tumors were harvested and processed as previously described (*45*). Briefly, tumors were mechanically dissociated and digested with collagenase/hyaluronidase containing DNase I at 37 °C for 30–45 minutes, lysed with ACK lysis buffer, and filtered with 40 μm cell strainer. For flow cytometry staining, cells (1×10^6^) were incubated with mouse Fc receptor blocker (#NB309, Innovex) at 4 °C for 10 minutes, then stained with the appropriate antibodies to surface markers at 4 °C for 30 minutes in the dark, and/or fixed/permeabilized (Cytofix/Cytoperm™ Fixation/Permeabilization Kit, #554714, BD Biosciences) and stained with intracellular antibodies at 4 °C for 30 minutes. A Zombie Aqua Fixable Viability Kit (#423101, BioLegend) was used to differentiate dead and live cells. The stained populations were analyzed using a Fortessa flow cytometer (BD Biosciences) and FlowJo software (BD Biosciences). The following antibodies were from BioLegend: anti-mouse CD45-BV785 (#103149), CD3-Alexa Fluor 647 (#100209), CD3-APC/Cy7 (#100221), CD11b-APC (#101212), CD4-FITC (#100510), CD8a-PE (#100708), CD8a-APC/Cy7 (#100714), CD19-PE/Cy5 (#115509), CD11c-PE/Cy7 (#117318), CD11c-APC/Cy7 (#117324), Ly6C-APC/Cy7 (#128026), Ly6G-FITC (#127605), F4/80-PerCP/Cy5.5 (#123127), F4/80-Alexa Fluor 488 (#123120), F4/80-BV605 (#123133), CD40-PE/Cy7 (#124621), CD206-FITC (#141704), CD69-FITC (#104505), CD107a-PE (#121612), PD1-FITC (#135213), CD44-PerCP/Cy5.5 (#103032), CD62L-APC (#104412), H-2KbOVA-APC (#141605), anti-human EpCAM-Alexa Fluor 647 (#324212), CD45-BV510 (#368525), and CD68-APC (#333810). The other antibodies were: CD11b-PE (#557397, BD Biosciences), CD335-eFluor 450 (#48335180, Invitrogen), iNOS-PE (#12592082, Invitrogen), and CD103-FITC (#11103182, Invitrogen).

### Immunofluorescence staining

Tumors were fixed in 4% PFA, infiltrated with 30% sucrose in PBS, and embedded in OCT compound. The lungs were processed as above. Frozen tissues were cryosectioned at the Histology Core at CSHL. OCT compound-embedded tissue sections were post-fixed with ice-cold methanol and acetone (1:1 ratio), rinsed with PBS, blocked with Fc receptor blocker (#NB309, Innovex), and incubated with 1 × blocking buffer (5% goat serum [#X0907, Dako], 2.5% BSA, 0.1% Triton X-100 in PBS) (*38*). Tumor sections were then incubated with fluorochrome-conjugated antibody (1:100, Alexa Fluor 488 anti-mouse F4/80, #123120, BioLegend; or PE anti-mouse CD40, #124610, BioLegend; or PE anti-iNOS, #12592082, Invitrogen; or FITC anti-mouse CD8a, #100706, BioLegend) in 0.5 × blocking buffer overnight at 4 °C. Lung sections were incubated with anti-PyMT antigen antibody (1:100, #ab15085, Abcam) overnight. The second day, tissue sections were counterstained with DAPI and mounted with mounting media (#17985-16, Electron Microscopy Sciences), while lung sections were incubated with Alexa Fluor 488 goat anti-Rat secondary antibody (1:1500, #A11006, Invitrogen) for 45 minutes, then incubated with Alexa Fluor 647 anti-mouse CD4 (1:100, #100424, BioLegend) and PE anti-mouse CD8a (1:100, #100708, BioLegend) overnight. The third day, lung sections were counterstained with DAPI and mounted with mounting media. Images were obtained with a Leica SP8 confocal microscope. The positive area of F4/80, CD8a, CD40, or iNOS in tumor sections was analyzed with ImageJ.

For the tumor re-challenge assay, lung sections were stained with anti-ECFP antibody (1:100, #AHP2986, Bio-Rad) and Alexa Fluor 488 goat anti-rabbit secondary antibody (1:1500, #A11070, Invitrogen). Images were scanned using an Aperio ScanScope® FL System (Leica Biosystems), and ECFP^+^ lung metastasis was determined using Aperio eSlide Capture Devices software (Leica Biosystems).

### In vitro T cell activation experiments

CD8^+^ T cells were isolated from the spleens of transplanted PyMT tumor-bearing mice with a CD8a^+^ T Cell Isolation Kit (#130-104-075, Miltenyi Biotec), according to the manufacturer’s instructions. Macrophages were differentiated from the bone marrow of these mice. The bone marrow was cultured for six days with 10 ng/ml mouse M-CSF (#CYT-439, ProSpec) on Not TC-treated culture dishes (#430597, Corning). The attached cells at the end of this culture period are unpolarized macrophages (*62*). CD8^+^ T cells were cultured with or without macrophages under different experimental conditions, as indicated below and in the figure legends, in RPMI-1640 with 20% FBS, 200 μM L-glutamine, 100 μM sodium pyruvate, and 50 μM 2-mercaptoethanol (all of these reagents in the T cell culture medium were from Gibco). Macrophages were seeded 2 hours before T cells to allow for their attachment, and MPLA (100 ng/ml, #tlrl-mpls, InvivoGen), mouse IFNγ (33 ng/ml, #485-MI/CF, R&D Systems), or the two together were added to the culture system immediately after the seeding of T cells. The activation of T cells was identified by CD69 expression using flow cytometry, similar to the experiments described above using cells isolated from tumors. To block the cytokines that activate CD8^+^ T cells, neutralizing antibodies against IL-12 (1 μg/ml, #AF-419-NA, R&D Systems), TNFα (5 μg/ml, #MAB4101, R&D Systems), or CXCL9 (6 μg/ml, #AF-492-NA, R&D Systems) were added to the co-culture system 30 minutes before MPLA with IFNγ.

For the antigen-presenting assay, CD8^+^ T cells were isolated from the spleens of OT1 mice (Jackson Laboratory) with a CD8a^+^ T Cell Isolation Kit (#130-104-075, Miltenyi Biotec). Macrophages were derived from the bone marrow of PyMT tumor-bearing mice as above and treated with MPLA (100 ng/ml, #tlrl-mpls, InvivoGen), IFNγ (33 ng/ml, #485-MI/CF, R&D Systems), or the two together for 16 hours, then incubated with OVA-Q4H7 peptide (10^−9^ M, #AS-64405, AnaSpec) for 2 hours. The pre-treated macrophages were washed with PBS and finally added to CD8^+^ T cells. CD69 expression of CD8^+^ T cells was identified by flow cytometry.

### Immune cell depletion/neutralization experiments

Anti-CD8 (#BE0004-1, BioXCell) and anti-CD4 (#BE0003-1) antibodies, or anti-CSF1R (#BE0213) antibody, were injected intraperitoneally one day before treatment with MPLA+IFNγ. For the first depletion, 0.5 mg of anti-CD4 and 0.5 mg of anti-CD8, or 1 mg of anti-CSF1R were injected per mouse. For the subsequent depletions, 0.2 mg of anti-CD4 and 0.2 mg of anti-CD8, or 0.5 mg of anti-CSF1R were injected per mouse.

### Statistics

Prism software (version 8.4.1, GraphPad) was used to analyze the data. In survival experiments, Kaplan-Meier survival curves were plotted and log rank test was performed. In all other experiments with only two groups, the unpaired t-test (equal variances) or unpaired Welch t-test (unequal variances) was used. One-way ANOVA (with equal variances) or Welch’s ANOVA (unequal variances) was performed for experiments with more than two groups. Bartlett’s test or Brown-Forsythe test was used to assess equality of variances among groups. When the ANOVA was significant, group means were compared as indicated in the figures and p-values adjusted for multiple comparisons were obtained using the Holm-Sidak procedure (equal variances) or Dunnett T3 procedure (unequal variances). All graphs present the mean, the error bars represent the standard deviation (s.d.), and p-values are presented in the figures. All p-values<0.05 were considered statistically significant.

## List of supplementary materials

### Supplementary materials and methods

#### Supplementary figures

Fig. S1. INOS activity is required for the tumoricidal effects of tumor-associated macrophages (TAMs) induced by MPLA with IFNγ.

Fig. S2. Administration of MPLA with IFNγ inhibits breast tumor growth in mice.

Fig. S3. MPLA with IFNγ enhances secretion of chemokines and stimulates type I interferon signaling in breast tumors.

Fig. S4. Gating strategy for flow cytometry of immune cells.

Fig. S5. MPLA with IFNγ induces recruitment of breast tumor-infiltrating leukocytes.

Fig. S6. Both macrophages and T cells are required for the anti-tumor effect of MPLA with IFNγ treatment.

Fig. S7. Intraperitoneal administration of MPLA with IFNγ suppresses ovarian cancer metastasis in mice.

#### Supplementary tables

Table S1. MPLA with IFNγ suppresses metastasis in ovarian tumor-bearing mice.

Table S2. The sequences of PCR primers.

#### Supplementary movies

Movie S1. Un-activated breast tumor-derived macrophages do not kill cancer cells.

Movie S2. MPLA with IFNγ activates breast tumor-derived macrophages to kill cancer cells.

#### Supplementary data file

Data file S1. List of upregulated and downregulated genes in the MPLA+IFNγ, MPLA and IFNγ treatment groups.

## Acknowledgments

We thank the patients participating in the research, as well as the staff of the Clinical Trial Office of New York University (NYU) Langone’s Perlmutter Cancer Center and the Center for Biospecimen Research and Development at NYU Langone Health for supporting this research.

## Author contributions

L.S., T.K., B.L., S.A., and M.E. designed the experiments; L.S., T.K., B.L., A.S.A., X.H., and D.N. performed the experiments; S.A. and I.A.M. provided essential reagents; P.G. verified all the statistical analyses; L.S. and M.E. wrote the manuscript; and T.K., B.L., A.S.A., D.S., I.A.M., X.H., D.N. and S.A. provided feedback on the manuscript.

## Data and materials availability

RNA sequencing data have been deposited in GSE146211.

