## Supplementary materials, methods and figures for "Activating a collaborative innate-adaptive immune response to control breast and ovarian cancer metastasis"

**Affiliations:**

**Supplementary Materials**

**Supplementary materials and methods**

**List of supplementary figures**

Fig. S1. INOS activity is required for the tumoricidal effects of tumor-associated macrophages (TAMs) induced by MPLA with IFNγ.

Fig. S2. Administration of MPLA with IFNγ inhibits breast tumor growth in mice.

Fig. S3. MPLA with IFNγ enhances secretion of chemokines and stimulates type I interferon signaling in breast tumors.

Fig. S4. Gating strategy for flow cytometry of immune cells.

Fig. S5. MPLA with IFNγ induces recruitment of breast tumor-infiltrating leukocytes.

Fig. S6. Both macrophages and T cells are required for the anti-tumor effect of MPLA with IFNγ treatment.

Fig. S7. Intraperitoneal administration of MPLA with IFNγ suppresses ovarian cancer metastasis in mice.

**List of supplementary tables**

Table S1. MPLA with IFNγ suppresses metastasis in ovarian tumor-bearing mice.

Table S2. The sequences of PCR primers.

**List of supplementary movies**

Movie S1. Un-activated breast tumor-derived macrophages do not kill cancer cells.

Movie S2. MPLA with IFNγ activates breast tumor-derived macrophages to kill cancer cells.

**List of supplementary data file**

Data file S1. List of upregulated and downregulated genes in the MPLA+IFNγ, MPLA and IFNγ treatment groups.

### Supplementary materials and methods

#### *MTS assay*

PyMT tumor cells were seeded to a 96-well plate ($5\times10^3$/well) and treated with LPS (100 ng/ml, #L4391, Sigma Aldrich), MPLA (100 ng/ml, #tlrl-mpls, InvivoGen), or polyI:C (1 µg/ml, #tlrl-picw, InvivoGen) with or without mouse IFNγ (33 ng/ml, #485-MI/CF, R&D) for 48 hours. Next, 20 µl CellTiter 96® AQueous One Solution reagent (#G3582, Promega) was added to 100 µl medium and incubated for 2 hours at 37 °C. Absorbance at 490 nm was measured using a SpectraMax MiniMax 300 Imaging Cytometer (Molecular Devices).

#### *Western blot analysis*

Macrophages were lysed by RIPA lysis buffer (#89990, Thermo Fisher Scientific) with protease and phosphatase inhibitor (#78440, Thermo Fisher Scientific). Protein extracts were resolved through 8% SDS-PAGE, transferred to polyvinylidene difluoride (PVDF) membrane (#1620177, Bio-Rad), probed with antibody against iNOS (1:1000, #610431, BD Biosciences), CD206 (1:1000, #AF2535, R&D) or β-actin (1:2000, #sc-47778, Santa Cruz Technology), and then with fluorescent secondary antibody (1:2000, #926-32210, LI-COR). Binding was visualized by an Odyssey® Classic Imaging System (LI-COR).

#### *ELISA for NETs*

Plasma samples were obtained by cardiac blood collection using a syringe with a 25 G needle containing 25 µl sodium citrate solution (#C3821, Sigma). Whole blood was centrifuged at 1,300 x g for 10 minutes at 4 °C, and the top plasma layer was collected. As in a previous study (*38*), 96-well Enzyme ImmunoAssay/Radio Immuno-Assay (EIA/RIA) plates (#3590, Costar) were coated overnight at 4 °C with an anti-elastase antibody (1:250, #sc-9521, Santa Cruz Biotechnology) in 15 mM $Na_2CO_3$ and 35 mM $NaHCO_3$ at pH 9.6. The next day, the wells were washed three times with PBS, blocked in 5% BSA for 2 hours at room temperature, and washed three times with PBS. Then, 50 µl plasma samples were added to the wells and incubated for 2 hours at room temperature on a shaker, and the plates were washed three times with a wash buffer (1% BSA, 0.05% Tween 20 in PBS). Next, anti-DNA-peroxidase conjugated antibody (1:50, #11774425001, Roche) in 1% BSA in PBS was added to the wells for 2 hours at room temperature, and the wells were washed five times with wash buffer before the addition of 2,2'-azino-bis(3-ethylbenzothiazoline-6-sulphonic acid) (ABTS, #37615, Thermo Fisher Scientific). Optical density was read 40 minutes later at 405 nm using a SpectraMax MiniMax 300 Imaging Cytometer (Molecular Devices).

***ELISA of cytokines***

The samples were collected as for analysis by cytokine array. CXCL9 and CCL5 secreted by tumors were assayed using a DuoSet ELISA (#DY492/#DY478, R&D Systems) and ancillary reagent kit (#DY008, R&D Systems) according to the manufacturer's guidelines. Absorbance at 450 nm was measured using a SpectraMax MiniMax 300 Imaging Cytometer (Molecular Devices).

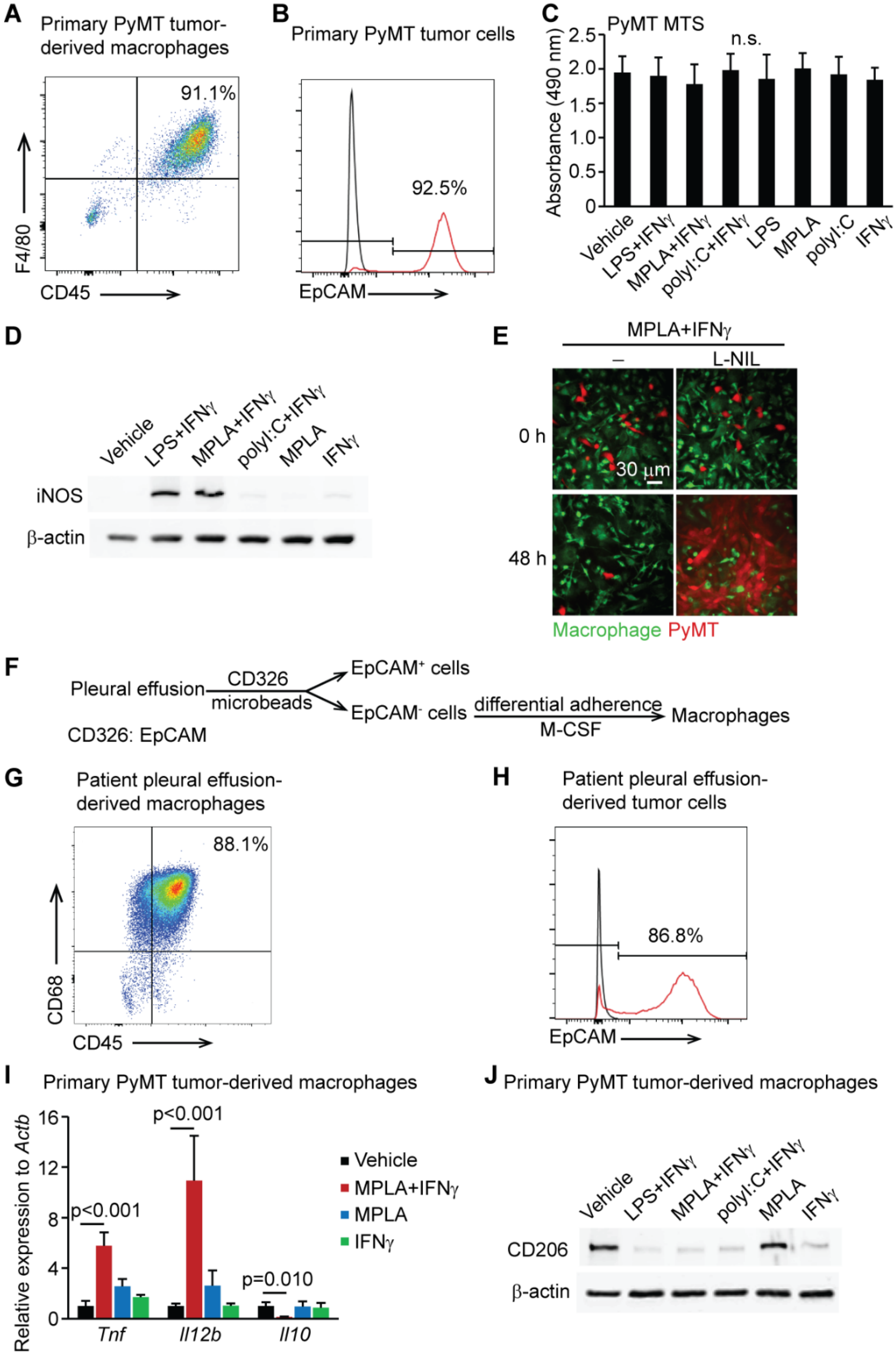


**Fig. S1. INOS activity is required for the tumoricidal effects of tumor-associated macrophages (TAMs) induced by MPLA with IFNγ.** (**A**) Purities of macrophages ($CD45^{+}F4/80^{+}$) and (**B**) cancer cells ($EpCAM^{+}$) from the primary PyMT tumors were identified by flow cytometry. (**C**) MTS assay of PyMT cancer cells. Cancer cells were isolated from the primary tumors of MMTV-PyMT mice and treated with indicated agents for 48 hours before measuring MTS activity. Graph depicts the average results of three independent experiments. One-way ANOVA was performed (mean±s.d.). n.s.: not significant. (**D**) INOS expression by macrophages was determined by Western blot. Macrophages were isolated from the primary tumors of MMTV-PyMT mice and treated with indicated agents for 36 hours before the assay. (**E**) Inhibition of iNOS activity blocked macrophages' tumoricidal activities. Macrophages and PyMT cancer cells were isolated from the primary tumors of MMTV-PyMT mice and co-cultured together with or without the iNOS activity inhibitor L-NIL during treatment and co-culturing. L-NIL: $N^6$-(1-Iminoethyl)-L-lysine hydrochloride. (**F**) Experimental procedure to isolate macrophages and cancer cells from the pleural effusions of breast cancer patients. (**G**) Purities of macrophages ($CD45^+CD68^+$) and (**H**) cancer cells ($EpCAM^+$) from the pleural effusions were identified by flow cytometry. Representative of three patient samples. (**I**) Expression of genes in primary PyMT tumor-derived macrophages was determined by RT-qPCR 24 hours after indicated treatments were initiated. Graph depicts the average results of three independent experiments. *Tnf* and *Il12b* (p40 subunit of Il12) are markers of tumoricidal macrophages. *Il10* is a marker of TAMs. The relative expression to *Actb* was normalized to vehicle, which was set to 1. One-way ANOVA was performed (mean±s.d.). (**J**) CD206 (a marker of TAMs) expression by primary PyMT tumor-derived macrophages was determined by Western blot. Macrophages were treated with indicated agents for 36 hours before the assay.

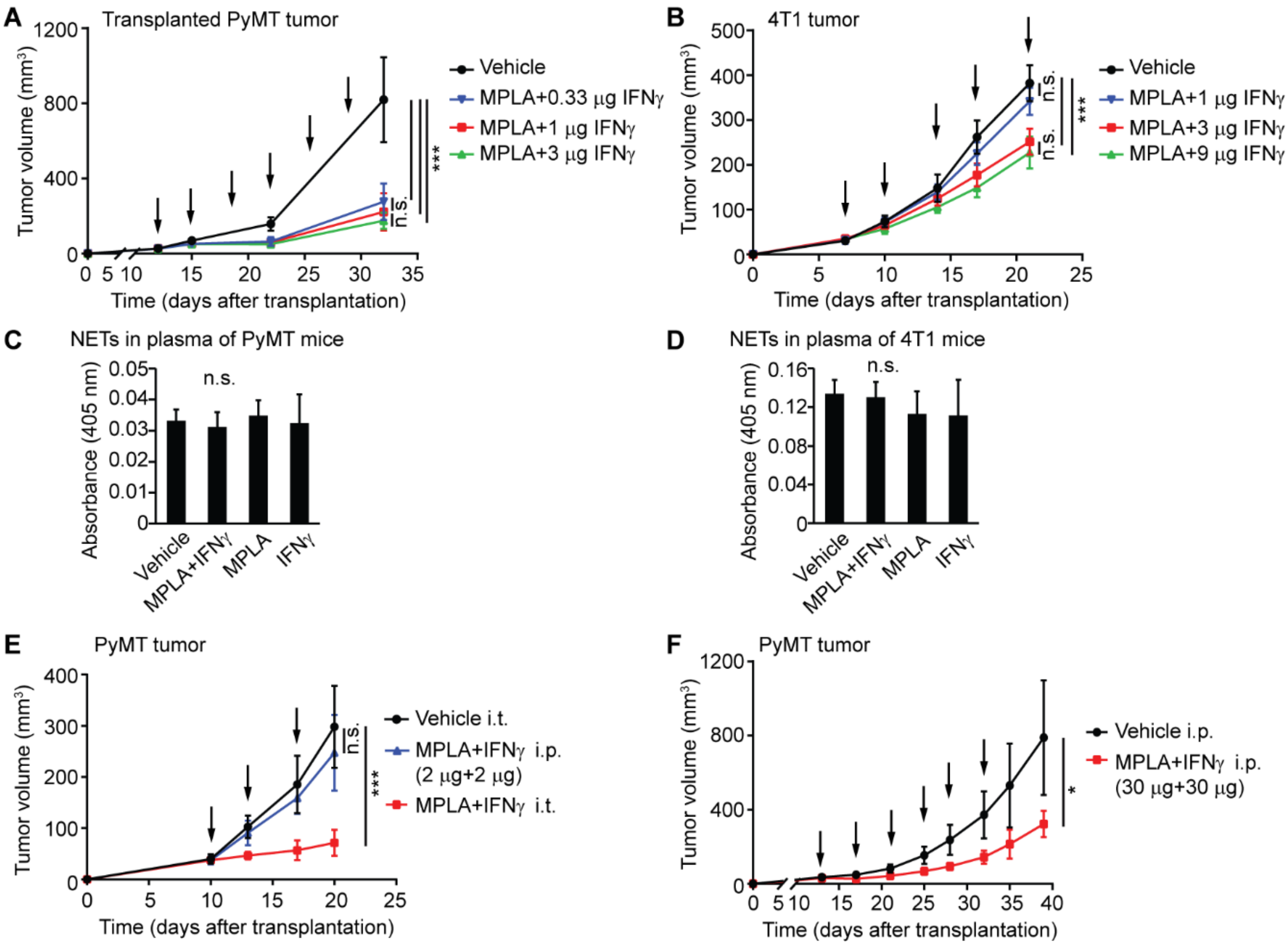


**Fig. S2. Administration of MPLA with IFNγ inhibits breast tumor growth in mice.** (**A**) Tumor growth curve of PyMT tumors treated with intratumoral injected MPLA and different doses of IFNγ. N=8 tumors per group. Note that the depicted vehicle and MPLA+1 µg IFNγ groups are the same as in Fig. 2A: the data are from one large experiment and shown as two separate graphs for clarity. Since the PyMT tumors in our model are derived from transplanted primary cells, the growth rate of vehicle-treated tumors shows variation between experiments, and we compared treatment only between groups of mice transplanted with the same primary cell population. (**B**) Tumor growth curve of 4T1 tumors treated with different doses of IFNγ. N=8 tumors per group. Note that the depicted vehicle and MPLA+3 µg IFNγ groups are the same as in Fig. 2C: the data are from one large experiment and shown as two separate graphs for clarity. (**C, D**) ELISA analysis of NETs in the plasma of PyMT tumor-bearing mice (**C**) or 4T1 tumor bearing-mice (**D**). N=5 mice per group; 1 µg MPLA, 1 µg mouse IFNγ, or the two together were used for PyMT tumors, while 1 μg MPLA, 3 μg mouse IFNγ, or the two together were used for 4T1 tumors. One-way ANOVA was performed for ELISA analysis in panels C and D (mean±s.d.). n.s.: not significant. (**E**) Tumor growth curve of PyMT tumors during treatment with low doses of MPLA with IFNγ. Arrows indicate time of treatments; 2 μg MPLA with 2 μg IFNγ per mouse was used. N=10 tumors per group. i.p.: intraperitoneal injection, i.t.: intratumoral injection. (**F**) Tumor growth curve of PyMT tumors during treatment with high doses of MPLA with IFNγ. Arrows indicate time of treatments; 30 μg MPLA with 30 μg IFNγ per mouse was used. N=5 tumors per group. One-way ANOVA (Fig. S2A, S2B, and S2E) or unpaired t-test (Fig. S2F) was performed for the tumor volume analysis at the end time point (mean±s.d.). Because the s.d. was significantly different between the groups (unequal group variances), Welch's ANOVA was used for Fig. S2A and S2E, and Welch's t-test was used for Fig. 2F. n.s.: not significant. *p<0.05, ***p<0.001.

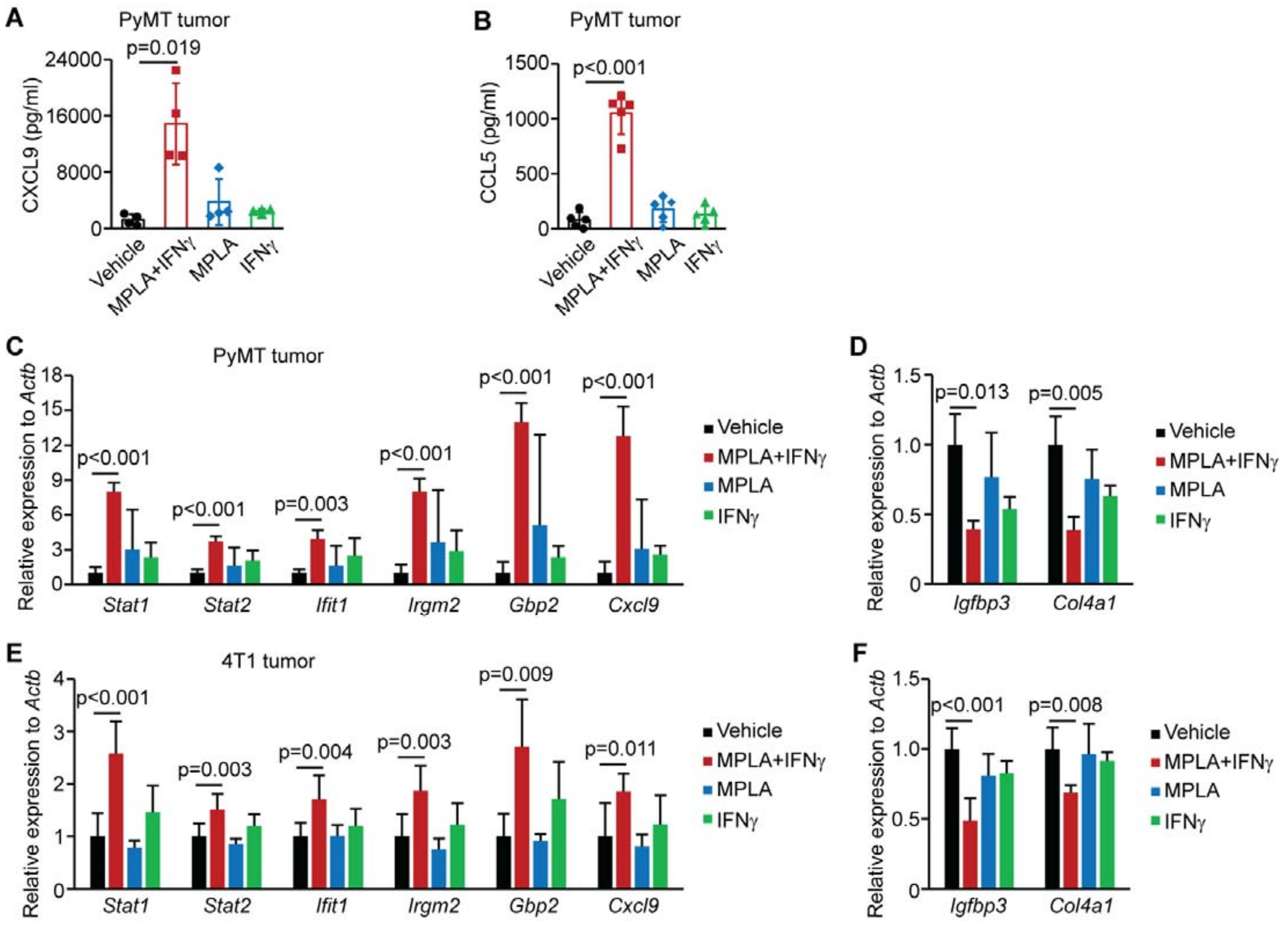


**Fig. S3. MPLA with IFNγ enhances secretion of chemokines and stimulates type I interferon signaling in breast tumors.** All PyMT and 4T1 tumors were treated and collected following the experimental design in Fig. 3A; 1 μg MPLA, 1 μg mouse IFNγ, or the two together were used for PyMT tumors, while 1 μg MPLA, 3 μg mouse IFNγ, or the two together were used for 4T1 tumors. (**A, B**) ELISA analysis of CXCL9 (**A**) and CCL5 (**B**) secreted by PyMT tumors after indicated treatments. (**C, D**) mRNA expression of indicated genes in PyMT tumors was determined by RT-qPCR. (**E, F**) mRNA expression of indicated genes in 4T1 tumors was determined by RT-qPCR. The genes belong to the type I interferon signaling pathway (C, E) or are ECM-associated genes (D, F). N=4–5 mice per group. The relative expression to *Actb* was normalized to vehicle, which was set to 1 (mean±s.d.). For each gene, one-way ANOVA was performed to compare expression between the different treatment groups, and due to unequal group variances, Welch's ANOVA was used for Fig. S3A, S3C (*Stat1*, *Stat2*, *Irgm2*, *Gbp2*, *Cxcl9*), S3D, S3E (*Gbp2*) and S3F (*Col4a1*).

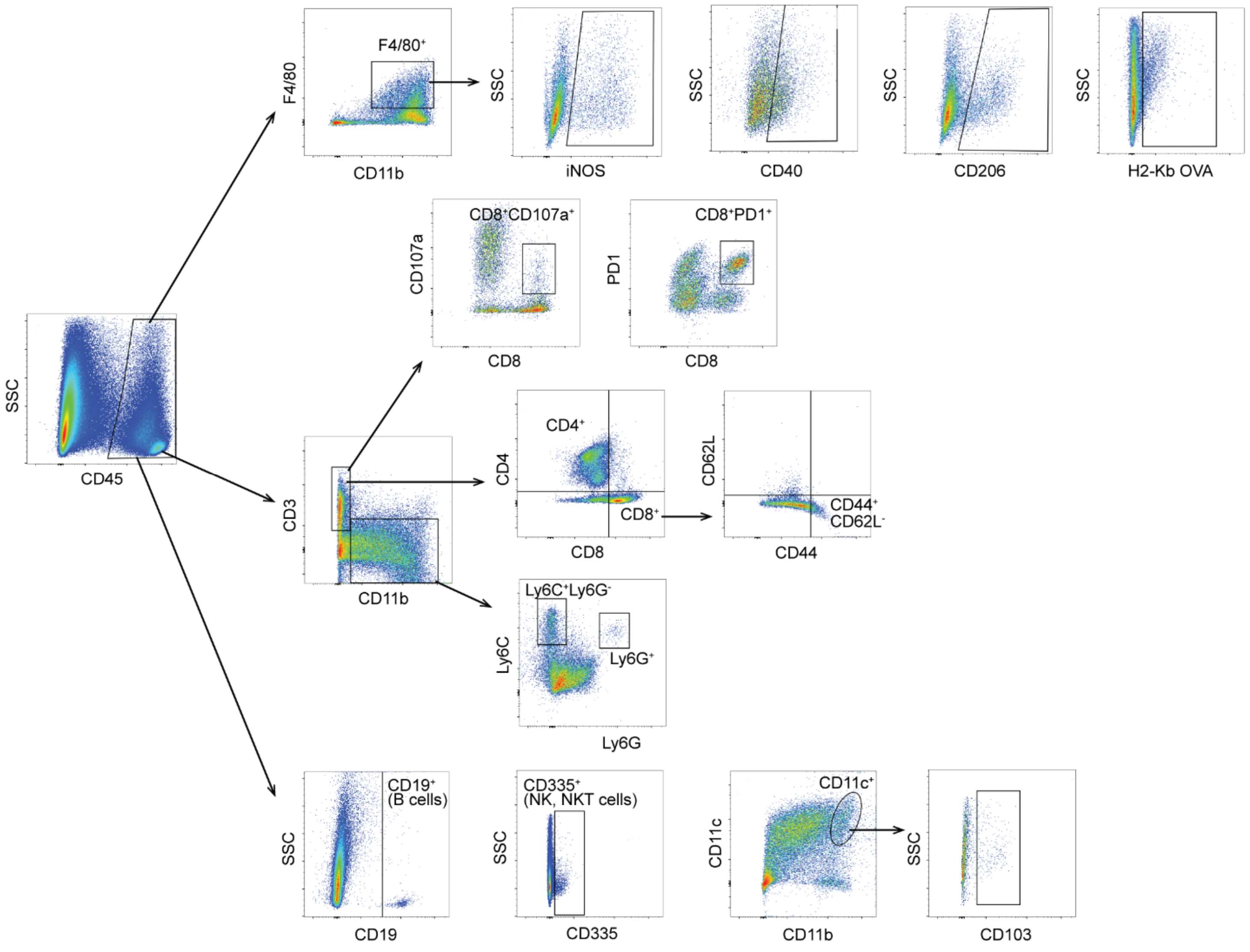


**Fig. S4. Gating strategy for flow cytometry of immune cells.** Two days after the 6$^{th}$ intratumoral injection, tumors were processed into single cells and flow cytometry was performed to determine the extent of immune cell infiltration into tumors. For flow cytometry analysis, single cells from tumors were gated on live CD45$^{+}$ cells and were further analyzed using the indicated markers to characterize the infiltration and activation of macrophages and T cells. SSC: side scatter.

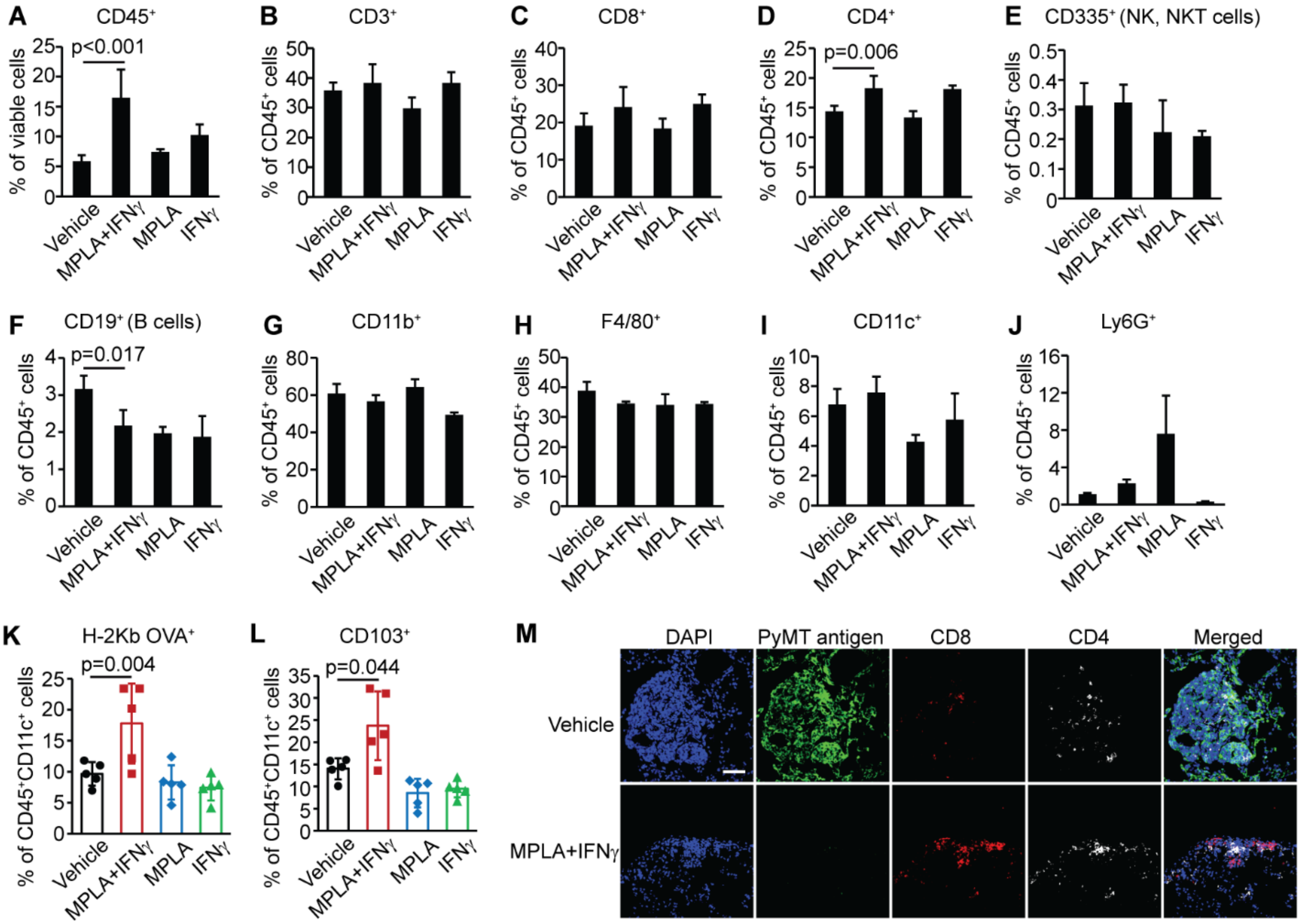


**Fig. S5. MPLA with IFNγ induces recruitment of breast tumor-infiltrating leukocytes.** (**A–J**) Infiltration of different immune cells in PyMT tumors was identified by flow cytometry using the strategy outlined in Fig. S4. N=3–6 mice per group. NK cells: natural killer cells, NKT cells: natural killer T cells. (**K–L**) The antigen-presenting activity of dendritic cells (H2-Kb OVA$^+$, **K**) and the proportion of CD103$^+$ CD11c$^{High}$CD11b$^{High}$ cells (**L**) in OVA$^+$ PyMT tumors were identified by flow cytometry. N=5 mice per group. In total, $2\times10^5$ cancer cells isolated from primary tumors of MMTV-PyMT-chOVA mice were transplanted into C57BL/6 mice to obtain these OVA$^+$ tumors. One-way ANOVA was performed for panels A–L (mean±s.d.), and due to unequal group variances, Welch's ANOVA was used for Fig. S5L. (**M**) T cell infiltration into the lungs of tumor-bearing mice was identified by IF staining. Scale bar: 50 μm. For all figure panels, the analysis was performed 2 days after the last intratumoral injection.

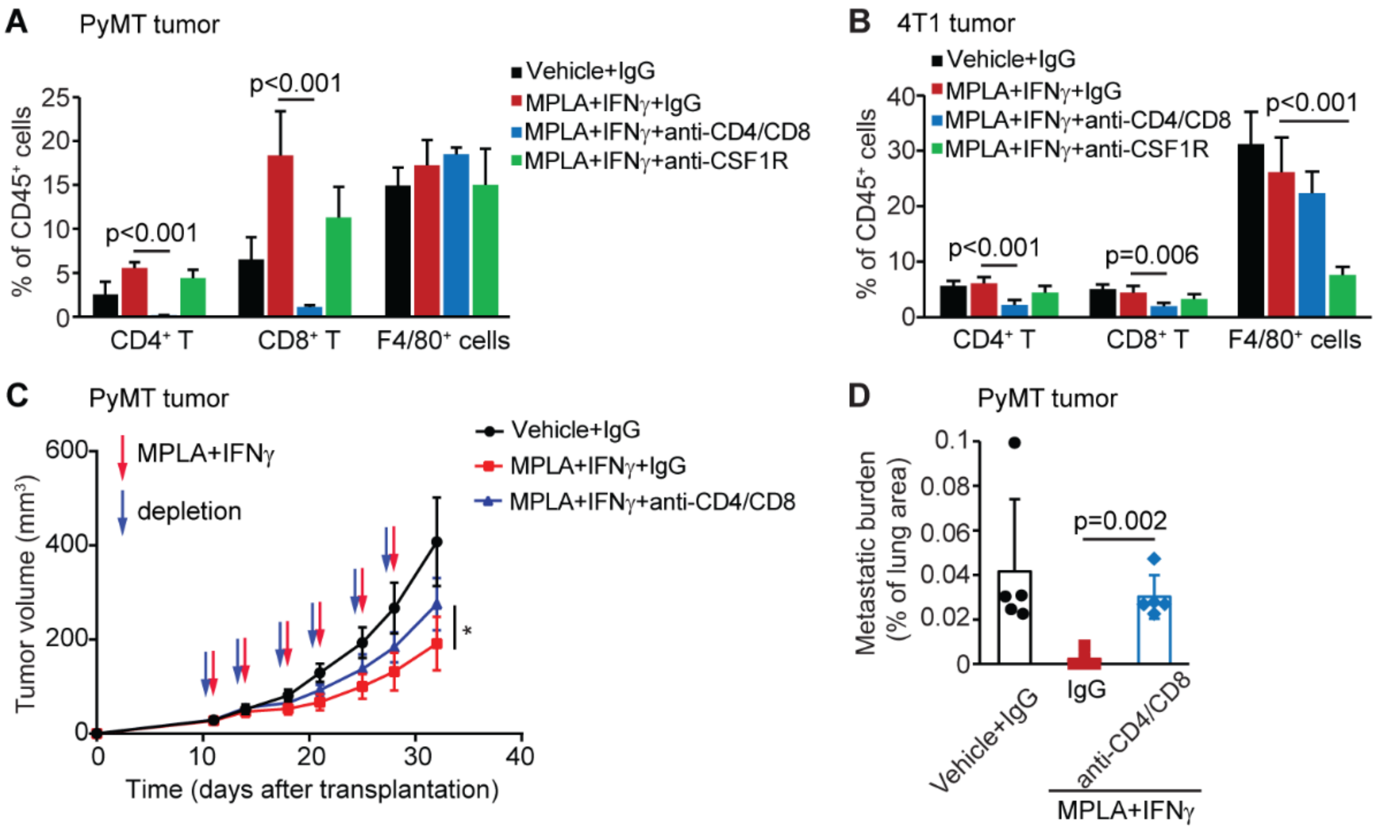


**Fig. S6. Both macrophages and T cells are required for the anti-tumor effect of MPLA with IFNγ treatment.** (**A, B**) Transplanted PyMT tumors (**A**) or 4T1 tumors (**B**) were removed, and flow cytometry was used to assay the depletion of T cells or macrophages in the tumor tissue. N=3–5 mice per group. One-way ANOVA was performed (mean±s.d.). (**C**) Tumor growth curves of PyMT tumors after T cell depletion. N=10 tumors per group. One-way ANOVA was performed for the tumor volume analysis at the end time point (mean±s.d.). *p<0.05. (**D**) Lung metastatic burden of PyMT tumor-bearing mice after T cell depletion. Lung tissues were collected three days after last MPLA+IFNγ treatment. N=5 mice per group. Welch's ANOVA was performed for Fig. S6D (mean±s.d.) because of unequal group variances.

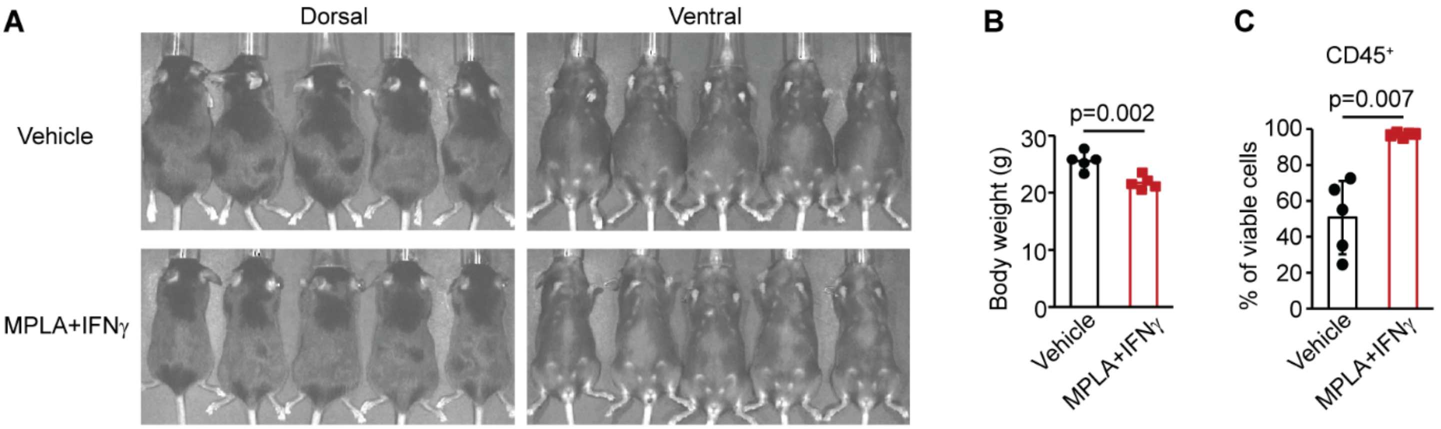


**Fig. S7. Intraperitoneal administration of MPLA with IFNγ suppresses ovarian cancer metastasis in mice.** (**A**) Photograph of the appearance of ID8-p53$^{-/-}$ tumor-bearing mice with or without MPLA+IFNγ treatment 48 days after tumor inoculation and two days after last treatment. (**B**) Body weight of the tumor-bearing mice with indicated treatment. (**C**) The percentage of immune cells (CD45$^{+}$) in ascites/peritoneal lavage was identified by flow cytometry. N=5 mice per group. Graphs are representative of at least two independent experiments. Unpaired t-test was performed (mean±s.d.), and due to unequal group variances, Welch's t-test was used for Fig. S7C.

**Table S1. MPLA with IFNγ suppresses metastasis in ovarian tumor-bearing mice.**

| Mouse group | Mouse number | peritoneum | pancreas | mesentery | diaphragm | kidney | spleen | adrenal gland |
|---|---|---|---|---|---|---|---|---|
| Vehicle | 1 | + | +++ | +++ | +++ | – | – | – |
| | 2 | +++ | +++ | +++ | +++ | ++ | +++ | +++ |
| | 3 | +++ | +++ | +++ | +++ | + | – | – |
| | 4 | +++ | +++ | +++ | +++ | – | – | – |
| | 5 | +++ | +++ | +++ | +++ | – | – | – |
| MPLA +IFNγ | 6 | + | + | – | – | – | – | – |
| | 7 | – | – | – | – | – | – | – |
| | 8 | – | + | – | + | – | – | – |
| | 9 | – | + | – | – | – | – | – |
| | 10 | ++ | + | – | – | – | – | – |

+, ++ and +++ indicate extent of macroscopic metastatic burden. –, not detected.

### Table S2. The sequences of PCR primers.

| Name | Sequence (5'–3') |
|---|---|
| CD40-S | tttggggtcaagcagattgc |
| CD40-AS | ttgcacaaccaggtctttgg |
| ACTB-S | tcgtgcgtgacattaaggag |
| ACTB-AS | ttgccaatggtgatgacctg |
| NOS2-S | ttgatgtccgaggcaaacag |
| NOS2-AS | acacgttcttggcatgcatg |
| TNF-S | tggctcagacatgttttccg |
| TNF-AS | gacagttggtcaccaaatcagc |
| CD206-S | cctggaaaaagctgtgtgtcac |
| CD206-AS | agtggtgttgcccttttgc |
| IL12B-S | atgccgttcacaagctcaag |
| IL12B-AS | atggcttcagctgcaagttc |
| IL10-S | acatcaaggcgcatgtgaac |
| IL10-AS | acggccttgctcttgttttc |
| Stat1-S | ggcctctcattgtcaccgaa |
| Stat1-AS | tgaatgtgatggccccttcc |
| Stat2-S | agaagtcctgcattggagcc |
| Stat2-AS | cttgttgccctttcctgcac |
| Ifit1-S | gtttgcatggctgaagtgca |
| Ifit1-AS | ttgggatggaattgcctgct |
| Irgm2-S | tgtcatcaagtaccacggcc |
| Irgm2-AS | gggggagaggggtgttatct |
| Gbp2-S | gctttaatgagccgcgactg |
| Gbp2-AS | cctccagcaagtctgagcaa |
| Cxcl9-S | aacctgcctagatccggact |
| Cxcl9-AS | cctgggatttggtgacgtga |
| Igfbp3-S | tctgggagcccataaggaca |
| Igfbp3-AS | ttgggcgtgtctgcagttat |
| Col4a1-S | aacaacgtctgcaacttcgc |
| Col4a1-AS | gcagaggcgagcatcatagt |
| Actb-S | aagtgtgacgttgacatccg |
| Actb-As | tctgcatcctgtcagcaatg |
| Tnf-S | tcgtagcaaaccaccaagtg |
| Tnf-AS | tttgagatccatgccgttgg |
| Il12b-S | ttgttcgaatccagcgcaag |
| Il12b-AS | acattcccgcctttgcattg |
| Il10-S | aaacaaaggaccagctggac |
| Il10-AS | ttccgataaggcttggcaac |

S: sense, AS: anti-sense

**Supplementary Movie Legends:**

**Movie S1. Un-activated breast tumor-derived macrophages do not kill cancer cells.** Vehicle-treated breast tumor-derived macrophages did not kill PyMT cancer cells. Macrophages and PyMT tumor cells were isolated from the primary tumors of MMTV-PyMT mice, and macrophages were stained with CellTracker™ Deep Red Dye, while tumor cells were stained with CellTracker™ CMFDA Dye. Deep Red dye is falsely colored green for clarity. Time indicated is time after imaging was initiated.

**Movie S2. MPLA with IFNγ activates breast tumor-derived macrophages to kill cancer cells.** Breast tumor-derived macrophages that were treated with MPLA+IFNγ killed PyMT cancer cells in 48 hours. Macrophages and PyMT tumor cells were isolated from the primary tumors of MMTV-PyMT mice, and macrophages were stained with CellTracker™ Deep Red Dye, while tumor cells were stained with CellTracker™ CMFDA Dye. Deep Red dye is falsely colored green for clarity. MPLA (100 ng/ml) with mouse IFNγ (33 ng/ml) was added before imaging was initiated. Time indicated is time after imaging was initiated.

**Supplementary Data File description:**

**Data file S1. List of upregulated and downregulated genes in the MPLA+IFNγ, MPLA and IFNγ treatment groups.** RNA-seq data were collected following the experimental design in Fig. 3A. After the combined MPLA+INFγ treatment, 179 genes were upregulated and 36 genes were downregulated in PyMT tumors, compared to vehicle. After treatment with MPLA alone, 20 genes were upregulated and 5 genes were downregulated. After treatment with INFγ alone, 33 genes were upregulated and 15 genes were downregulated.
